# MSD2, an apoplastic Mn-SOD, contributes to root skotomorphogenic growth by modulating ROS distribution in Arabidopsis

**DOI:** 10.1101/2021.12.01.470564

**Authors:** Huize Chen, Jinsu Lee, Jung-Min Lee, Minsoo Han, Aurélia Emonet, Jiyoun Lee, Xingtian Jia, Yuree Lee

**Affiliations:** Higher Education Key Laboratory of Plant Molecular and Environmental Stress Response in Shanxi Province, Shanxi Normal University, Taiyuan, 030000, Shanxi, P. R. China; Research Institute of Basic Sciences, Seoul National University, Seoul, 08826, Republic of Korea; Research Center for Plant Plasticity, Seoul National University, Seoul 08826, Republic of Korea; School of Biological Sciences, Seoul National University, Seoul 08826, Republic of Korea; Department of Plant Molecular Biology, University of Lausanne, Biophore Building, UNIL-Sorge, 1015 Lausanne, Switzerland; Department of New Biology, DGIST, Daegu 42988, Republic of Korea; Plant Genomics and Breeding Institute, Seoul National University, Seoul 08826, Republic of Korea; Max Planck Institute for Plant Breeding Research, Carl-von-Linné-Weg 10, Cologne, 50829, Germany

**Keywords:** Superoxide dismutase, ROS metabolism, skotomorphogenesis, light response, root growth

## Abstract

Reactive oxygen species (ROS) play essential roles as a second messenger in various physiological processes in plants. Due to their oxidative nature, ROS can also be harmful. Thus, the generation and homeostasis of ROS are tightly controlled by multiple enzymes. Membrane-localized NADPH oxidases are well known to generate ROS during developmental and stress responses, but the metabolic pathways of the superoxide (O2·^−^) generated by them in the apoplast are poorly understood, and the identity of the apoplastic superoxide dismutase (SOD) is unknown in Arabidopsis. Here, we show that a putative manganese SOD, MSD2 is secreted and possesses a SOD activity that can be inhibited by nitration at tyrosine 68. The expression of *MSD2* in roots is light condition-dependent, suggesting that MSD2 may act on ROS metabolism in roots during the light-to-dark transition. Root architecture is governed by ROS distribution that exhibits opposite gradient of H_2_O_2_ and O2·^−^, which is indeed altered in etiolated *msd2* mutants and accompanied by changes in the onset of differentiation. These results provide a missing link in our understanding of ROS metabolism and suggest that MSD2 plays a role in root skotomorphogenesis by regulating ROS distribution, thereby playing a pivotal role in plant growth and development.

## 1. Introduction

Plant growth is accompanied by continuous biochemical reactions in response to environmental stimuli, resulting in redox imbalances. Reactive oxygen species (ROS), unavoidable metabolic by-products, play important signaling roles as mediators of metabolic and signaling processes [1]. ROS are highly reactive and oxidize biological molecules, causing cellular damage and affecting regulation of gene expression [2]. ROS accumulation and distribution are delicately regulated by complex reactions catalyzed by enzymes including plasma membrane– bound NADPH-oxidases (respiratory burst oxidase homolog, RBOH), which produce the superoxide (O2·^−^) in the apoplast; and superoxide dismutases (SODs), which metabolize O2·^−^ to hydrogen peroxide (H_2_O_2_), the most stable and least cytotoxic ROS [3]. In addition to serving as antioxidants under different stresses [4, 5], SODs are also involved in developmental processes during plant growth [6, 7].

SODs can be classified into one of four types based on their metal cofactors: Cu/Zn-SOD (copper/zinc), Fe-SOD (iron), Mn-SOD (manganese), and Ni-SOD (nickel), which are not present in eukaryotes other than the green alga *Ostreococcus tauri* [8-10]. The Arabidopsis (*Arabidopsis thaliana*) genome is thought to encode three Cu/Zn-SOD (CSD1, CSD2, and CSD3), three Fe-SOD (FSD1, FSD2, and FSD3), and one Mn-SOD (MSD1) enzymes [9]. They localize in different cellular compartments: CSD1 and FSD1 localize in the cytosol, CSD2 and FSD1-3 in the chloroplast, CSD3 in peroxisomes, and MSD1 in the mitochondria [5, 11, 12]. FSD1 also localizes in the nucleus to some extent [11]. SODs secreted to the extracellular space have not yet been reported in Arabidopsis.

SOD activity is generally a reflection of gene expression and protein abundance. However, post-translational modifications (PTMs) are also critical for SOD activity [13]. Phosphorylation of Ser/Thr residues or Tyr residues is a typical post-translational protein modification. Phosphorylation on the Ser-38 residue of Cu/Zn-SOD Sod1p in budding yeast (*Saccharomyces cerevisiae*) blocks enzymatic activity under hypoxia [14]. The cytoplasmic Mn-SOD of the bacterium *Listeria monocytogenes* is phosphorylated on Ser and Thr residues, resulting in a decrease in enzymatic activity when bacteria reach stationary phase [15]. Nitration is another extensively studied modification of SODs. Human Cu/Zn-SOD and Mn-SOD lose most of their enzymatic activity by peroxynitrite treatment [16, 17]. In Arabidopsis, peroxynitrite also inhibits MSD1, CSD3, and FSD3 activity via tyrosine nitration [18].

The Arabidopsis Mn-SOD, MSD1, localizes in mitochondria and plays pivotal roles during female gametogenesis and fertilization by regulating local ROS bursts [19]. A recent update to The Arabidopsis Information Resource (TAIR) database proposed a new SOD member, MSD2, which is predicted to be an Mn^2+^-bound SOD. Unlike other SODs, MSD2 presents a secretory peptide and thus has the potential to be secreted into the apoplast [1]. MSD2 may therefore act as a component of the ROS metabolic pathways generated in the apoplast by NADPH oxidase, which has not been elucidated so far. Here, we investigated the possible function of MSD2 by characterizing its SOD activity, protein modifications, and subcellular localization, as well as the expression pattern of the encoding gene. Our results revealed that MSD2 has SOD activity and is secreted into the apoplast. *MSD2* expression levels were low in the light but highly induced in the roots of etiolated seedlings. Phenotypic analysis of *msd2* mutant roots indicated that MSD2 modulates ROS distribution, affecting root architecture in etiolated seedlings. These results not only elucidate the molecular mechanism of ROS metabolism during root development and morphogenesis, but also provide insight that can be broadly applied to extracellular ROS-mediated signaling pathways.

## 2. Materials and methods

### 2.1. Plant materials and growth conditions

The Columbia ecotype of *Arabidopsis thaliana* was used. T-DNA insertion mutants *msd2-1* (GABI_100H05) and *msd2-2* (SM_3_35975) were obtained from the ABRC, and *cop1-4* [20] was kindly provided by Dr. Young Hun Song (Seoul National University). Arabidopsis seeds were surface sterilized, sown on half-strength Murashige and Skoog (MS) plates (4.4 g/L MS salt, 1% [w/v] sucrose, pH 5.7, and 0.8% [w/v] agar), and stratified for 3 d at 4°C in the dark. The plates were then exposed to white light for 6 h to promote germination and subsequently kept either in darkness (covered with foil paper) or in continuous light in growth chambers with 60% relative humidity at 22°C (16-h-light/8-h-dark photoperiod, 22°C day/18°C night regime).

### 2.2. Plasmid Construction and Plant Transformation

Gateway cloning technology (Invitrogen) was used to generate all constructs: *pMSD2:nlsGFP-GUS, pUBQ10:MSD2-mCherry*, and *pUBQ10:MSD2*Δ*SP-mCherry*. Mutagenesis of MSD2/Y68F was conducted with the Fast Site-Directed Mutagenesis Kit (cat. n. KM101) from TianGen. The constructs were introduced into plants by the floral dip method [21]. In brief, Arabidopsis flowers were dipped into a solution containing Agrobacterium (*Agrobacterium tumefaciens*, GV3101), 5% (w/v) sucrose, and 0.05% (v/v) surfactant Silwet L-77 for 2 to 3 s with gentle agitation. Dipped plants were kept for 16 to 24 h in high humidity conditions. Transformed plants were selected using appropriate selection with antibiotics or herbicide. Plasmid constructs and primer details are given in Table S4.

### 2.3. Phylogenetic tree analysis and structure prediction

The protein sequences of SODs were obtained from the National Center for Biotechnological Information (NCBI) database by BLASTP. Protein sequence details are given in Table S1. Multiple protein alignment was performed with Clustal Omega, and the phylogenetic tree was constructed with MEGA X software using the neighbor-joining method [22]. For the modeling of the MSD2 structure, the MSD2 protein sequence was aligned to that of Arabidopsis MSD1 (PDB code: 4C7U) as template to model MSD2 using SWISS-Model [23]. Predictions of PTM sites for MSD2 were performed with the online tool MusiteDeep for phosphorylation and ubiquitination [24] and with the software GPS-YNO2 for Tyr-nitration [25]. PlantCARE was used for *in silico* analysis of *MSD2* promoter sequences [26], and TBtools was used for visualization [27].

### 2.4. Protein extraction and immunoblotting

Five days after germination (DAG), Arabidopsis seedlings were harvested and ground with a TissueLyser II (QIAGEN) in liquid nitrogen, and total protein was extracted using lysis buffer (10 mM Tris-Cl pH 7.5, 150 mM NaCl, 0.5 mM EDTA, 0.5% [v/v] NP 40, 0.09% [w/v] sodium azide). After centrifugation at 13,000 *g* for 20 min at 4°C, MSD2-mCherry was immunoprecipitated with RFP-trap agarose beads (Chromotek, cat. n. rtak-20). Immunoblotting of MSD2-mCherry was performed after native-PAGE or SDS-PAGE (12.5% separating gel and 4% stacking gel) with anti-mCherry primary antibody (Invitrogen, cat. n. M11217) diluted to 1:5,000 in Tris-buffered saline with 0.1% (v/v) Tween-20 (TBST) and 5% (w/v) low-fat dry milk. Tyr-nitration of MSD2 was detected after SDS-PAGE with anti-Nitrotyrosine primary antibody (Invitrogen, cat. n. A-21285) diluted to 1:3,000 in TBST with 3% (w/v) low-fat dry milk. Phosphorylation of MSD2 was detected by anti-phospho Ser/Thr and anti-phospho Tyr primary antibodies (Abcam, cat. n. ab117253 and RM111) both diluted to 1:3,000 in TBST with 3% (w/v) low-fat dry milk. The secondary antibodies were goat anti-rat (Invitrogen, cat. n. A10549) and goat anti-rabbit (Invitrogen, cat. n. G-21234). Signal intensity for each protein band was calculated by ImageJ software (National Institutes of Health, https://imagej.nih.gov/ij/) [28]. Contrast and brightness were adjusted in the same manner for all images. All analyses were performed in three biological replicates, and statistical significance was evaluated using a one-way analysis of variance (ANOVA) test.

### 2.5. SOD activity assay

For in-gel staining of SOD activity, native-PAGE was performed on a 15% gel at 4L; the samples were loaded without addition of SDS or heating [29]. After native-PAGE, the gel was incubated first with Nitroblue tetrazolium (NBT) (0.1% [w/v] in distilled water) solution, followed by incubation in riboflavin (28 μM in phosphate-buffered saline [PBS]) solution, with washes in water in between each step. The gel was exposed to light with a white-light box for 20 min at room temperature to reveal the white SOD activity bands over the blue background. For inhibitor treatments, the gel was immersed in 8 mM H_2_O_2_ (Fe-SOD and Cu/Zn-SOD inhibitor) for 30 min with shaking at room temperature before SOD activity staining as above. For inhibition of Cu/ZnSOD, 8 mM KCN was added to the riboflavin solution during SOD activity staining in the dark. Peroxynitrite (100 or 200 μM) was added to the protein extract in the dark for 20 min at room temperature [18]. To determine the optimum pH of MSD2 activity, SOD activity was measured in the pH range of 3.0 to 10.0 using 50 mM sodium citrate (pH 3.0–8.0), Tris-HCl (pH 8.0–9.0), or glycine-NaOH (pH 9.0–11.0) buffers [30]. SOD isoenzyme activity was measured according to the manufacturer’s instructions in the SOD activity assay kit (Abcam, cat. n. ab65354).

### 2.6. Subcellular localization, GUS, and ROS staining

For propidium iodide (PI) staining, 5-DAG seedlings were incubated in the dark in 10 μg/mL PI (Invitrogen, cat. n. P4170) for 10 min and then rinsed twice with water and observed under a Zeiss LSM 700 confocal microscope (Zeiss, Germany)[31]. For the plasmolysis experiments, 5-DAG seedlings were treated with 0.8 M mannitol [32] with 0.1% (w/v) Calcofluor white (Sigma-Aldrich, cat. n. 18909) for 30 min. For GUS staining, 3-, 7-, and 11-DAG seedlings were incubated in GUS staining solution (3 mM potassium ferrocyanide, 3 mM potassium ferricyanide, 1 mM X-Gluc, 1 M NaH_2_PO_4_, 1 M Na_2_HPO_4_, 0.5 M EDTA, pH 8.0) at 37°C for 6 h. Seedlings were then rinsed with 70% (v/v) ethanol for at least 5 min, destained for chlorophyll in clearing solution (ethanol:acetic acid = 6:1, v/v), and then rinsed with 70% (v/v) ethanol and observed under a microscope (Leica, M205FA). To check ROS distribution, roots from seedlings grown on MS plates for 5 DAG were used. For staining hydrogen peroxide, seedlings were incubated in 5 μM 2-[6-(4′-hydroxy) phenoxy-3H-xanthen-3-on-9-yl] benzoic acid, hydroxyphenyl fluorescein (HPF) staining solution (in 1/2 MS, pH = 5.8) for 5 min at room temperature. For staining superoxide, seedlings were incubated in 5 μM dihydroethidium (DHE) staining solution (in 1/2 MS, pH = 5.8) for 30 min at 37°C. The roots were washed with 1/2 MS media and imaged by fluorescence microscopy (Leica, M205FA). The quantitative measurement of ROS staining intensity was performed as average pixel intensities using ImageJ software [28].

### 2.7. RT-qPCR analysis of gene expression

Seedlings were harvested at 7 DAG and snap-frozen in liquid nitrogen. Total RNA was extracted using an EasyPure Plant RNA Kit (Transgen, cat. n. ER301-01). The extracted RNA was treated with RNase-free DNase I. First-strand cDNA was synthesized by reverse transcription using TransScript One-Step RT-PCR SuperMix (Transgen, Cat. n. AT411-02), and the obtained cDNAs were used for gene expression analysis by qPCR. Each qPCR reaction contained 12.5 μL TransStart Tip Green qPCR SuperMix (Transgen, cat. n. AQ141-01), 0.1 μg cDNA, and 7.5 pmol of each gene-specific primer (Supplementary Table S1) in a final volume of 25 μL and was performed on a Bio-Rad CFX 96 real-time PCR system. *UBQ5* (At3g62250) was used as the reference gene, and relative gene expression levels were calculated using the 2^−ΔΔCT^ method. Three independent biological replicates were analyzed.

### 2.8. Confocal laser scanning microscopy

Transgenic seedlings expressing *mCherry-MSD2* were used for microscopy. Confocal laser scanning microscopy images were obtained using a Zeiss LSM 700 (with Zen SP3 Black edition). The excitation and detection windows were as follows: PI: 535 nm, 600–630 nm; mCherry: 560 nm, 600–630 nm; and GFP and HPF: 488 nm, 500–530 nm.

### 2.9. Statistical analysis

All statistical analyses were conducted in GraphPad Prism software v.8.0.0. One-way ANOVA was performed, and a two-sided *t*-test was subsequently used as a multiple-comparison test. Details about the statistical approaches used can be found in the figure legends. The data are presented as means ± standard deviation (SD) where indicated. Each experiment was repeated at least three times.

## 3. Results

### 3.1. Phylogenetic Analysis of Arabidopsis MSD2 in the plant Mn-SOD family

We built a phylogenetic tree based on protein sequences to investigate the relationship between the putative SOD AtMSD2 and SODs from various plant species. AtMSD2 was distantly related to other known Arabidopsis SODs (Fig. 1). AtMSD2 grouped with other MSDs in the MSD clade rather than in the AtCSD or AtFSD clade. The phylogenetic tree classified AtMSD2 and AtMSD1 into two distinct subclades, indicating that they are distantly related.

**Fig. 1.**
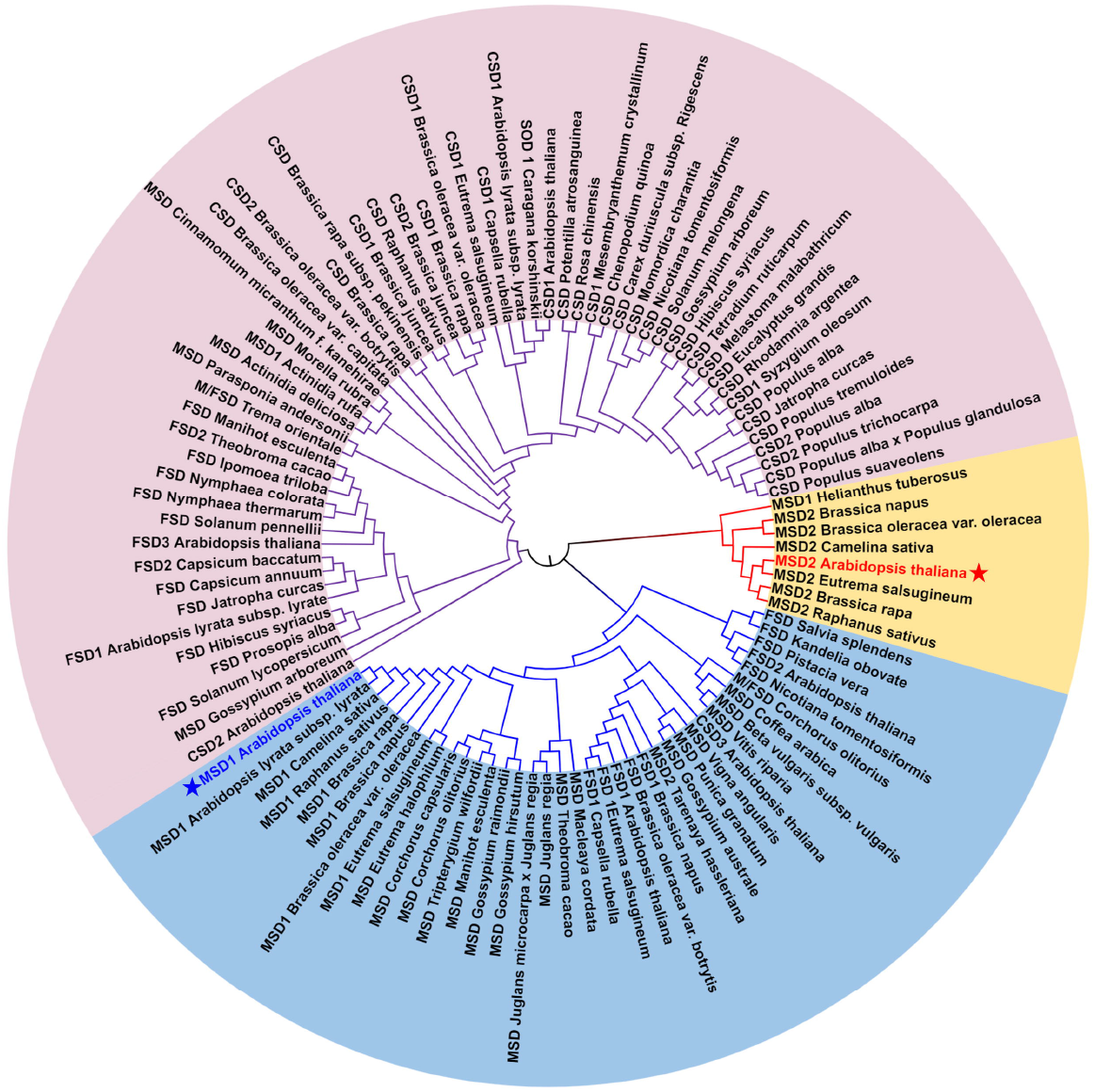
Phylogram of SODs from different plant species. Amino acid sequences of SODs from different plant species were obtained from the NCBI database by BLASTP. Multiple protein alignment was conducted with Clustal Omega, and the phylogenetic tree was constructed using the neighbor-joining method in MEGA X software. AtMSD1 and AtMSD2 are indicated in blue and red, respectively, and by stars. The SODs used for phylogenetic analysis are listed in Table S1.

### 3.2. Biochemical analysis of the SOD activity of MSD2

We performed in-gel SOD activity assays to assess whether MSD2 has SOD activity. After separating proteins by native-PAGE, we detected SOD activity by Nitroblue tetrazolium (NBT) and flavin staining. Compared to the wild type (WT), we observed additional SOD activity bands in transgenic seedlings overexpressing *MSD2-mCherry* (OE). Immunoblotting with an anti-mCherry antibody confirmed the new band seen on the native-PAGE gels is MSD2-mCherry (Fig. 2A). The SOD activity associated with MSD2-mCherry and MSD1 was not affected by incubation of the gels with potassium cyanide (KCN), an inhibitor of CSD activity, or H_2_O_2_, which inhibits FSDs and CSDs (Fig. 2B). We also tested the optimum pH for MSD2 activity on MSD2-mCherry immunoprecipitated from OE seedlings: MSD2 activity was strongest at pH 8.0 (Fig. 2C).

**Fig. 2.**
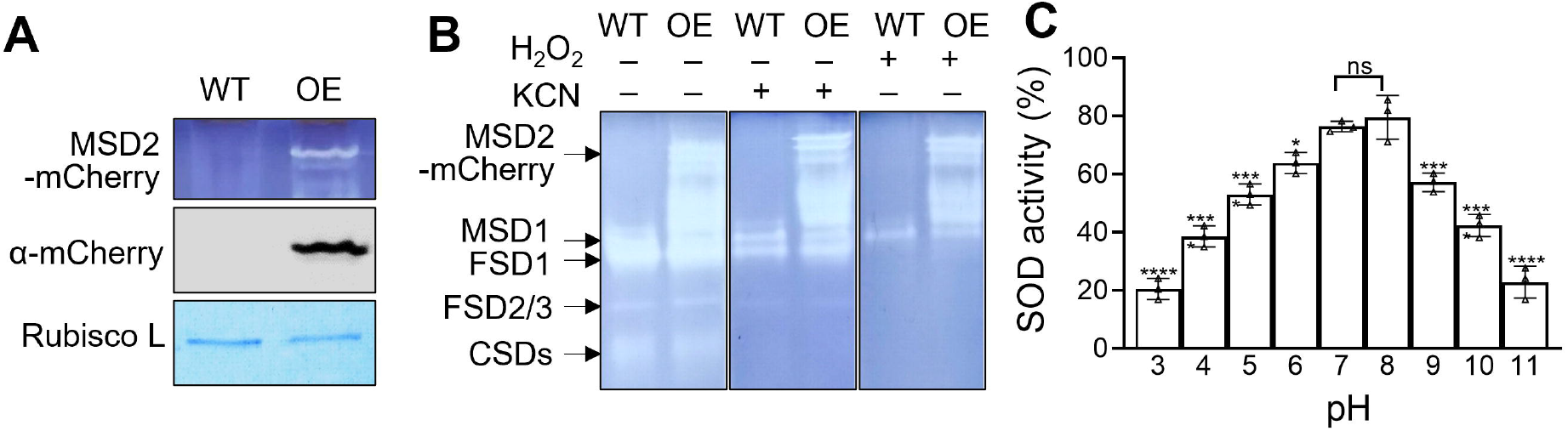
Characterization of MSD2 SOD activity. (A) Immunoblot analysis after native PAGE showing the presence of MSD2-mCherry in 14-day-old OE seedlings. WT, wild type; OE, *pUBQ10:MSD2-mCherry*. (B) Visualization of SOD isozymes by native PAGE in the indicated genotypes. KCN (inhibitor of CSD activity) and H_2_O_2_ (inhibitor for FSDs and CSDs) were used as specific SOD inhibitors. (C) Quantification of MSD2 SOD activity at various pH conditions *in vitro* after immunoprecipitation. Values represent means ± standard deviation (SD) of three independent experiments. Asterisks indicate significant differences (*t*-test, ** *P* < 0.01; *** *P* < 0.001; **** *P* < 0.0001); ns, non-significant.

### 3.3. Tyr-nitration induced by peroxynitrite inhibits MSD2 SOD activity

SOD activity is highly regulated through transcriptional regulation of the encoding gene as well as through PTM [13]. To elucidate the potential effects of PTM on MSD2, we searched the MSD2 protein sequence for putative PTM sites (Table S2), of which Tyr-68 (Y68) was identified as the most likely site for nitration (Table S3). To determine if Y68 is a target residue for nitration, we analyzed the activity center of MSD2. Modeling of the three-dimensional structure of MSD2 revealed that residues H60, H108, D197, and H201 form an active pocket around the manganese cation (Mn^2+^) (Fig. 3A). Alignment of MSD1 (PDB code: 4C7U) as a template showed that although the amino acid residues in the active pocket are different between MSD1 and MSD2, the underlying spatial structure is similar. In particular, Y68 in MSD2 completely overlapped at the same location with Y63 of MSD1, which is targeted by nitration [18] (Fig. 3B). The distance between the Y68 side chain and the Mn^2+^ within the active site decreased upon the addition of an – NO_2_ group at Y68 compared to the unmodified residue (Fig. 3C and D). Immunoblotting of MSD2-mCherry after immunoprecipitation with an anti-NO2-Tyr antibody indicated that MSD2-mCherry can undergo Tyr-nitration induced by peroxynitrite (Fig. 3E). Exogenous application of peroxynitrite inhibited MSD2 activity in a concentration-dependent manner, with 100 µM peroxynitrite lowering MSD2 activity by 20% and 200 µM peroxynitrite further decreasing MSD2 activity by 40% (Fig. 3F). Importantly, mutating Y68 to phenylalanine (F) greatly limited the inhibitory effects of peroxynitrite (Fig. 3E, F), suggesting that Y68 of MSD2 is the main target for nitration and inactivation of the enzyme. We also tested phosphorylation of immunoprecipitated MSD2 from OE seedlings by immunoblotting with anti-Ser/Thr/Tyr-phos antibodies but failed to detect any phosphorylation events (Fig. S1).

**Fig. 3.**
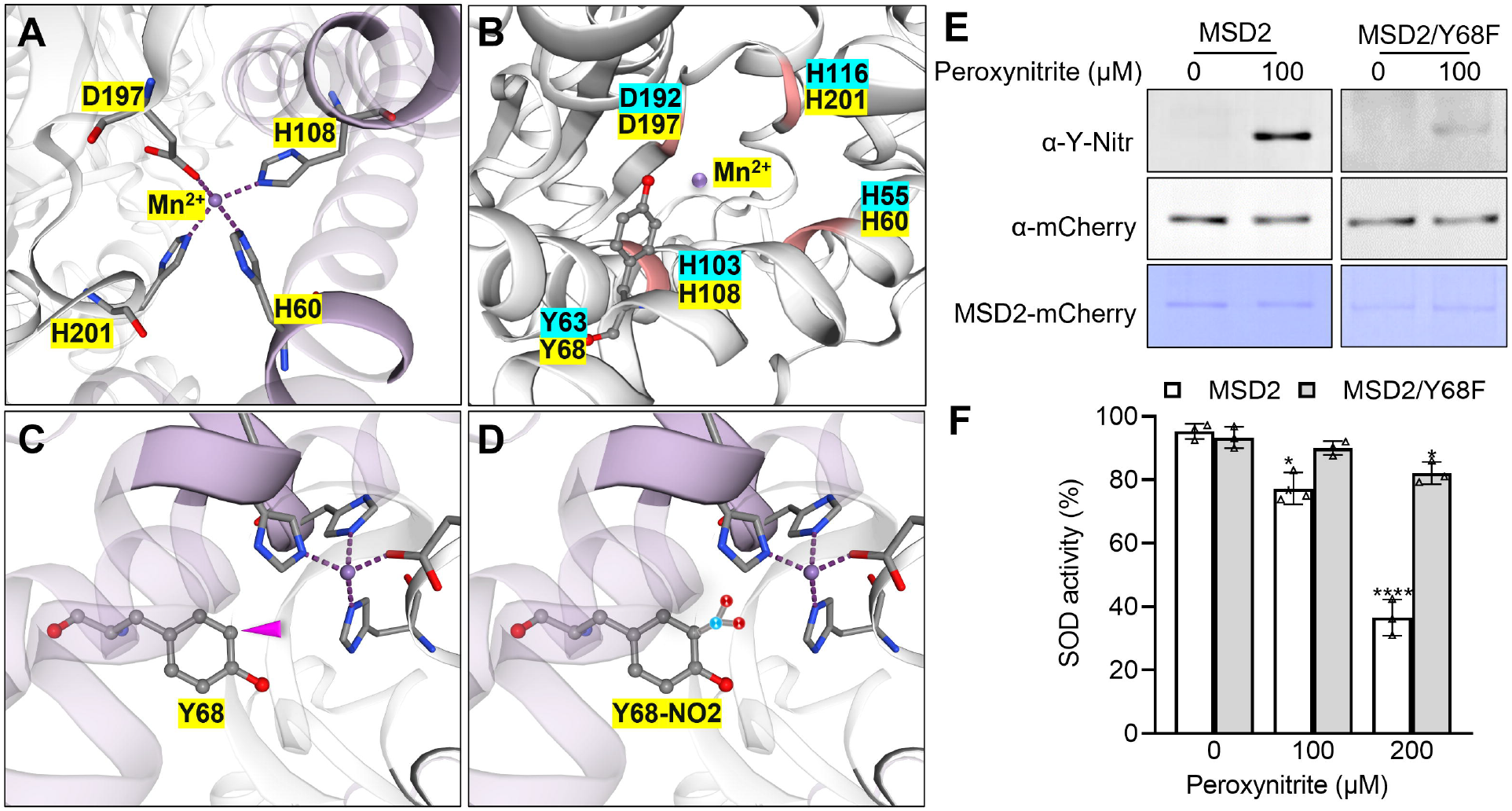
Structural prediction model of MSD2 nitration and effect of peroxynitrite on MSD2 activity. (A) The “active pocket” of MSD2, as generated with SWISS-MODEL using Arabidopsis MSD1 (PDB code: 4C7U) as guide. (B) Part of the structural model of MSD2 showing the nitration residues and the active pocket, aligned to MSD1. Residues with a blue background are from MSD1, and those with a yellow background are from MSD2. (C) Modeling of the substrate binding site of MSD2, modeled with unmodified Y68. The pink triangle indicates the –NO_2_ binding site. (D) Modeling of the substrate binding site shown with nitrated Y68. (E) Detection of nitrated Tyr residues by immunoblotting with anti-NO_2_-Tyr antibody on immunoprecipitates from *pUBQ10:MSD2-mCherry* and *pUBQ10:MSD2/Y68F-mCherry* seedling extracts. (F) Effect of peroxynitrite on SOD enzyme activity for MSD2 (white bars) and MSD2/Y68F (gray bars). Values are means ± SD from three independent experiments. Asterisks indicate significant differences (*t*-test, ** *P* < 0.01; *** *P* < 0.001; **** *P* < 0.0001).

### 3.4. Subcellular localization of MSD2 in Arabidopsis seedlings

MSD2 had a putative signal peptide in its N terminus and was predicted to be a secreted protein (Fig. S2). To assess the localization of MSD2 in plants, we examined the fluorescence signals in leaf epidermal cells of 5-DAG seedlings expressing *pUBQ10:MSD2-mCherry* or *pUBQ10:MSD2*Δ*SP-mCherry*, in which the potential signal peptide had been removed. We mainly detected MSD2-mCherry in the apoplast of epidermal cells. The fluorescence from mCherry remained associated with the cell wall after plasmolysis, clearly arguing against a plasma membrane localization for the fusion protein (Fig. 4A). In guard cells, we observed a strong signal in the vacuole, as well as a weak signal in the cell wall after plasmolysis, suggesting that some of it was secreted into the apoplast (Fig. 4B). Deleting the putative signal peptide abolished both the apoplast and vacuolar localization of MSD2-mCherry, resulting in the accumulation of MSD2ΔSP-mCherry in the cytoplasm (Fig. 4C, D).

**Fig. 4.**
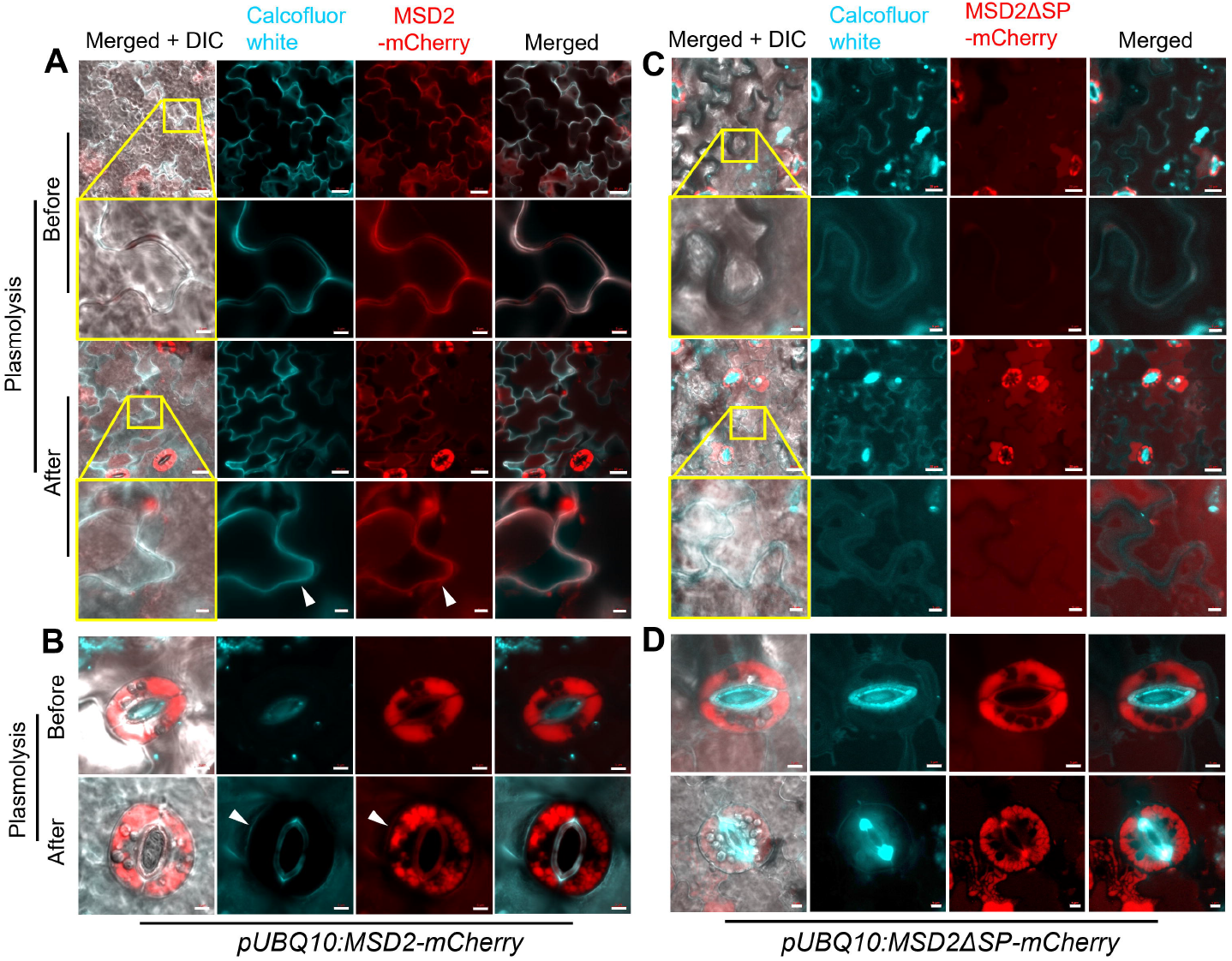
Subcellular localization of MSD2-mCherry and MSD2ΔSP-mCherry in Arabidopsis leaf epidermal cells. (A-D) Confocal microscopy images of Arabidopsis leaf epidermal cells expressing *pUBQ10:MSD2-mCherry* (A, B) or expressing *pUBQ10:MSD2*Δ*SP-mCherry* (C, D). Arrowheads indicate the cell wall. Calcofluor white was used to show the cell wall. Bars = 20 µm (A, C) and 5 µm (magnified view of A and C, and B and D).

### 3.5. *MSD2* expression in roots is regulated by light conditions

To characterize the expression of *MSD2* in Arabidopsis, we generated transgenic plants expressing *pMSD2:nlsGFP-GUS*. In 3-DAG seedlings, we observed strong GUS staining at the junction of the hypocotyl and the root (Fig. 5 A, B) and at the root tip (Fig. 5C). We confirmed the accumulation of the fusion protein at the root-shoot junction by confocal observations of the signal from the green fluorescent protein (GFP) moiety (Fig. 5D). We also detected GFP fluorescence in the vascular tissue of the root differentiation zone (Fig. 5E) and in root caps cells at the root tip (Fig. 5G).

**Fig. 5.**
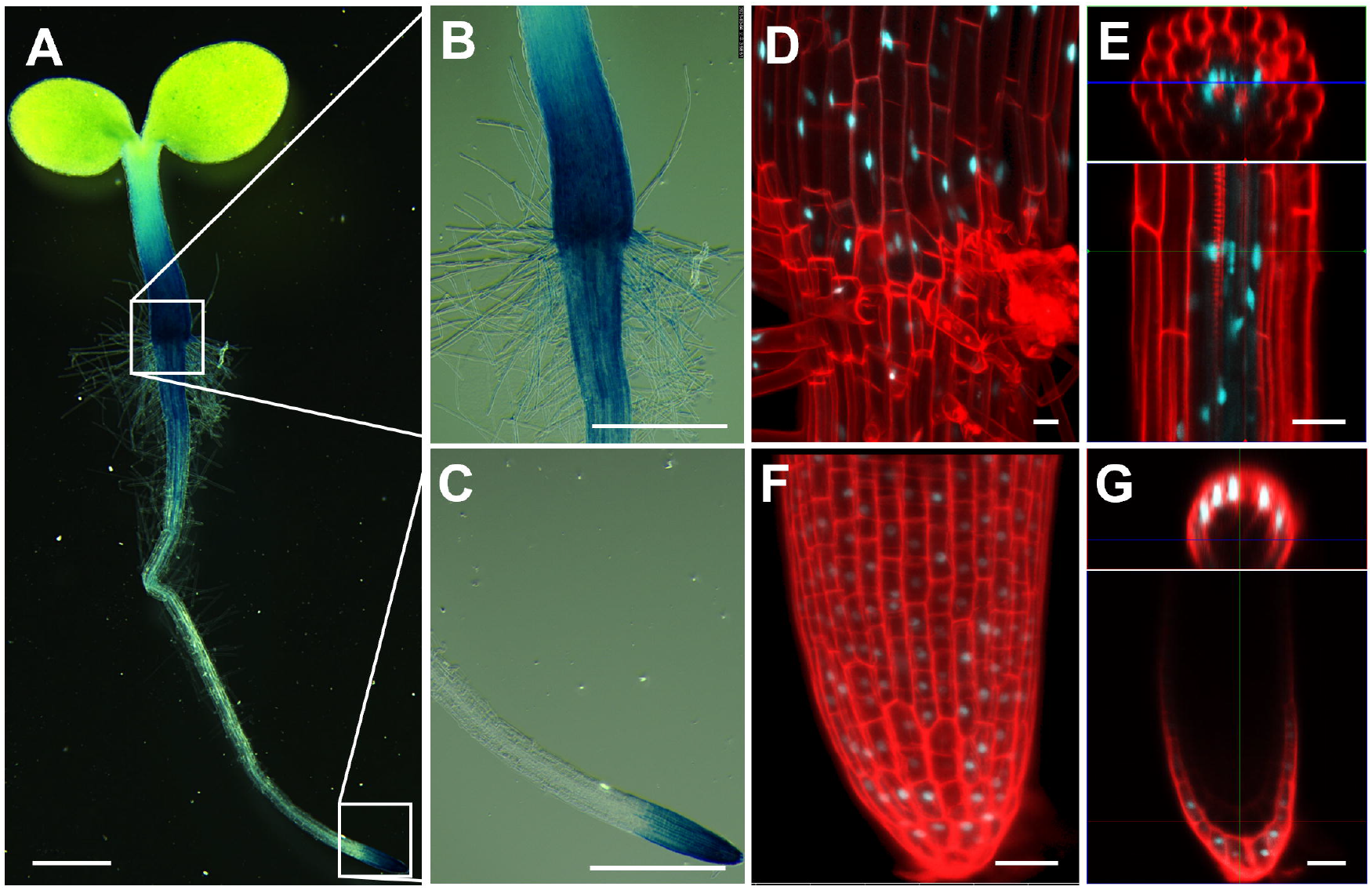
Promoter analysis of *MSD2* using the reporter line *pMSD2:nlsGFP-GUS*. (A-C) GUS staining pattern of a whole seedling. (B) and (C) are magnified views of the GUS staining pattern in the root-hypocotyl junction region and the root tip, indicated by white squares in (A). (D-F) Confocal microscopy images showing nlsGFP accumulation in the junction region (D), mature zone (E), and root tip (F, G). (G) is a cross-section image of nlsGFP signal in the root cap. Roots were stained with propidium iodide to visualize the cell wall (shown in red). nlsGFP is shown in cyan. Bars =1 mm (A), 500 µm (B-C), and 20 µm (D-G).

We analyzed the *cis*-elements of the *MSD2* promoter to explore the transcriptional regulation of *MSD2* expression using PlantCARE [26]. We determined that the *MSD2* promoter region contains *cis*-acting elements involved in abscisic acid signaling and responses to light (Fig. 6A). To investigate light-dependent changes in *MSD2* expression, we compared GUS patterns in seedlings grown in light or dark conditions. While the spatial GUS pattern did not change significantly when seedlings were exposed to light, we observed a dramatic change in signal intensity in seedlings grown in the dark (Fig. 6B-E). In the roots of 7-DAG seedlings, GUS signal was restricted to the vascular tissue, but staining intensity significantly increased in seedlings grown in the dark compared to light-grown seedlings (Fig. 6B, C). We also noticed higher *MSD2* expression in the lateral roots of 11-DAG plants grown in the dark relative to light-grown seedlings (Fig. 6D, E). We confirmed that *MSD2* expression is much higher in etiolated seedlings compared to light-grown seedlings by RT-qPCR (Fig. 6F). Photoreceptors are expressed in roots and contribute to regulating root morphogenesis via downstream signaling cascades that include the E3 ubiquitin ligase CONSTITUTIVE PHOTOMORPHOGENIC1 (COP1) and the transcription factor ELONGATED HYPOCOTYL5 (HY5) [33, 34]. As a major repressor, COP1 regulates the protein stability of photomorphogenesis-promoting transcription factors, including that of HY5 [35]. To test whether *MSD2* expression was under control of COP1, we measured *MSD2* expression in *cop1-4* mutants. *MSD2* expression was equally low in light-grown WT and *cop1-4* seedlings; by contrast, *MSD2* expression was much higher in *cop1-4* than in the WT in dark-grown seedlings (Fig. 6F), suggesting that COP1 affects *MSD2* expression in the dark.

**Fig. 6.**
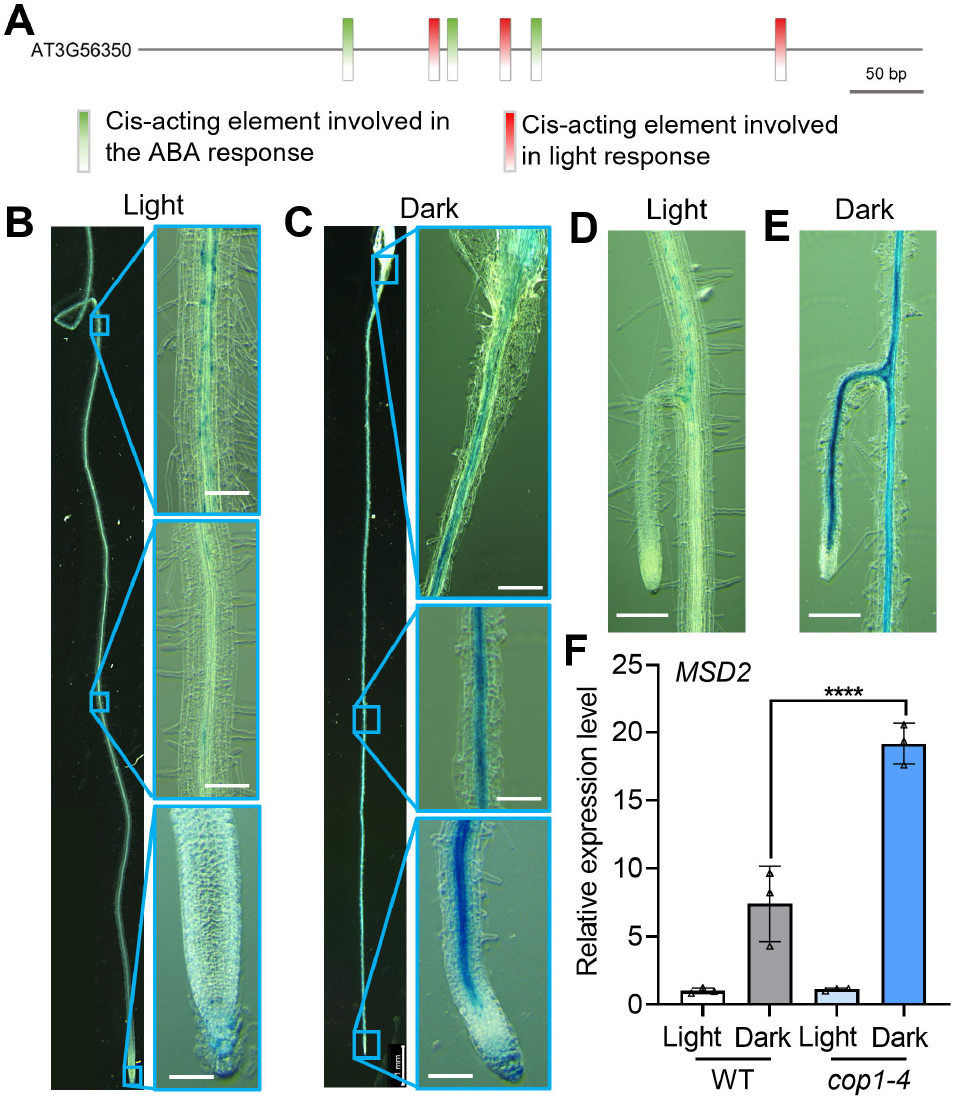
*MSD2* expression is regulated by light. (A) The *MSD2* promoter harbors *cis*-elements involved in light and abscisic acid (ABA) responses. (B, C) GUS staining in 7-DAG seedlings grown in the light (B) or in the dark (C). (D, E) GUS pattern in the lateral root of 11-DAG seedlings grown in the light (D) or in the dark (E). (F) RT-qPCR analysis of *MSD2* expression in the WT and *cop1-4* in light-grown and dark-grown seedlings. Expression levels were normalized to *UBQ5* (At3g62250) and expressed relative to light-grown WT. Bars = 50 bp (A) and 200 µm (B, C, D, E). Values are means ± SD of three independent experiments. Asterisks indicate significant differences (*t*-test, **** *P* < 0.0001).

### 3.6. MSD2 mediates skotomorphogenesis by modulating ROS distribution

Light conditions have a significant influence on root growth and architecture [36, 37]. To test whether MSD2 plays a role in regulating root morphogenesis, we obtained two independent T-DNA insertional mutants, *msd2-1* and *msd2-2*, in which the expression of *MSD2* was abolished (Fig. S3). We compared the root phenotypes of WT and *msd2-1* and *msd2-2* seedlings grown in the light or in the dark. The root length of light-grown seedlings was much longer than seedlings grown in the dark for both the WT and the *msd2* mutants. Under the dark condition, the root length in the *msd2-1* and *msd2-2* mutants was slightly longer than that of the WT, which was restored by introducing the *MSD2* gene in *msd2-1* mutants (Fig. 7A). Furthermore, we did notice a clear difference between the WT and mutants in the starting position of root hairs (Fig. 7B). In dark-grown seedlings, root hairs started closer to the root tip than in light-grown seedlings, but both *msd2-1* and *msd2-2* mutants exhibited a delay in the onset of root hair formation compared to the WT.

**Fig. 7.**
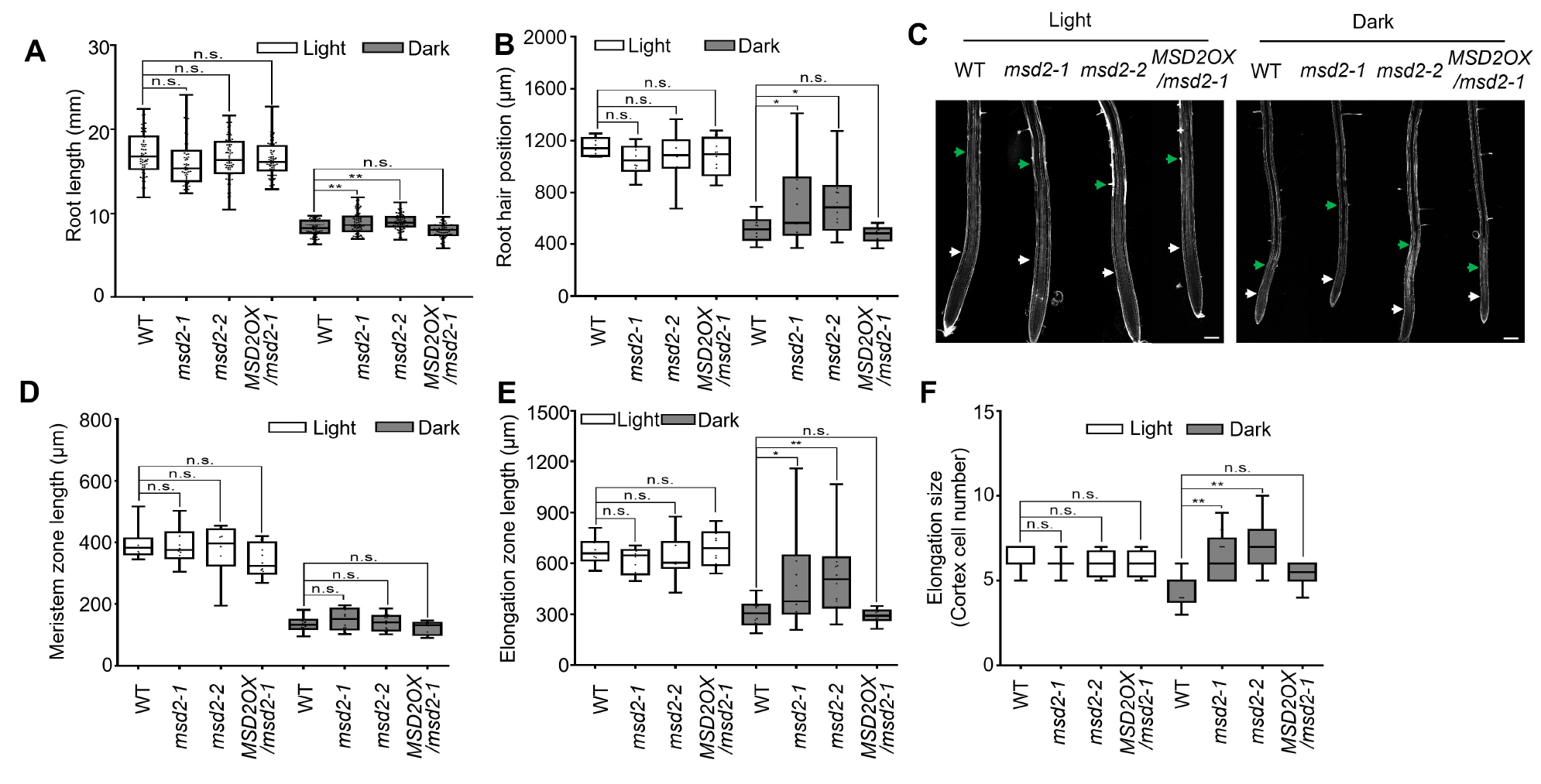
Root phenotypes of the *msd2* mutants. (A, B) Quantification of root length (A) and initiation sites of root hairs (B) of 5-DAG seedlings grown in the light or in the dark. (C) Confocal microscopy images of 5-DAG roots of the WT and the *msd2* mutant stained with propidium iodide. The end of the meristematic zone (white arrowheads) and the end of the elongation zone (green arrowheads) are indicated. (D-F) Mean length of the meristematic zone (D), the elongation zone (E), and mean cortex cell number (F) in the elongation zone of WT and *msd2* seedlings. *pUBQ10:MSD2-mCherry* was introduced in *msd2-1* mutants (*MSD2OX*/*msd2-1*). Bars = 100 µm. Values represent means ± SD of three independent experiments. Asterisks indicate significant differences (*t*-test, * *P* < 0.05, ** *P* < 0.01, *** *P* < 0.001).

ROS are known to play important roles in root growth [38]. In particular, the distribution of O2·^−^ and H_2_O_2_ are key to distinguishing the meristem zone from the differentiation zone [39, 40]. Since MSD2 possesses SOD activity, which would dismutate O2·^−^ to H_2_O_2_ (Fig. 2), and because *MSD2* expression was regulated by the light conditions, we speculated that this delay in root hair formation in dark-grown *msd2* seedlings might be related to the regulation of the transition from proliferation to differentiation influenced by O2·^−^ and H_2_O_2_ distribution. To test this possibility, we analyzed the size of the meristematic and elongation zones in WT and *msd2* mutant seedlings after staining the roots with propidium iodide (Fig. 7C). The size of the meristematic and elongation zones did not differ significantly between the WT and *msd2* in light-grown seedlings, but the elongation zone increased in *msd2-1* and *msd2-2* mutants compared to the WT in dark-grown seedlings (Fig. 7D, E). The increased size of the elongation zone in the *msd2* mutant appeared to be caused by an increase in cell number (Fig. 7F). These results suggest that *MSD2*, whose expression was induced in dark conditions, participates in skotomorphogenesis by limiting the elongation zone under dark conditions.

To test whether the effects of MSD2 on morphogenesis in dark conditions were related to the distribution of ROS, we determined the ROS distribution in roots using 3′-(*p*-hydroxyphenyl) fluorescein (HPF) to detect H_2_O_2_, and dihydroethidium (DHE) to detect O2·^−^ [40]. Consistent with previous studies [39, 40], we detected opposing gradients of HPF fluorescence and DHE fluorescence in the WT. HPF fluorescence was strong in the elongation zone and weak in the meristem zone (Fig. 8A, B), whereas DHE fluorescence was strong in the meristem zone and weak in the elongation zone (Fig. 8C, D). No significant differences could be found in light-grown *msd2* mutant seedlings, whereas in dark-grown seedlings, we observed a difference in ROS distribution between the WT and the *msd2* mutants (Fig. 8). HPF fluorescence in dark-grown *msd2* mutants was very low, and there was no clear gradient between meristem zone and elongation zone (Fig. 8B), but the DHE fluorescence in the elongation zone was stronger than that in the WT (Fig. 8D), showing that the opposing gradients of H_2_O_2_ and O2·^−^ were altered in *msd2* mutants. These results suggest that MSD2 contributes to ROS distribution in dark-grown seedlings, thereby controlling the transition between proliferation and differentiation.

**Fig. 8.**
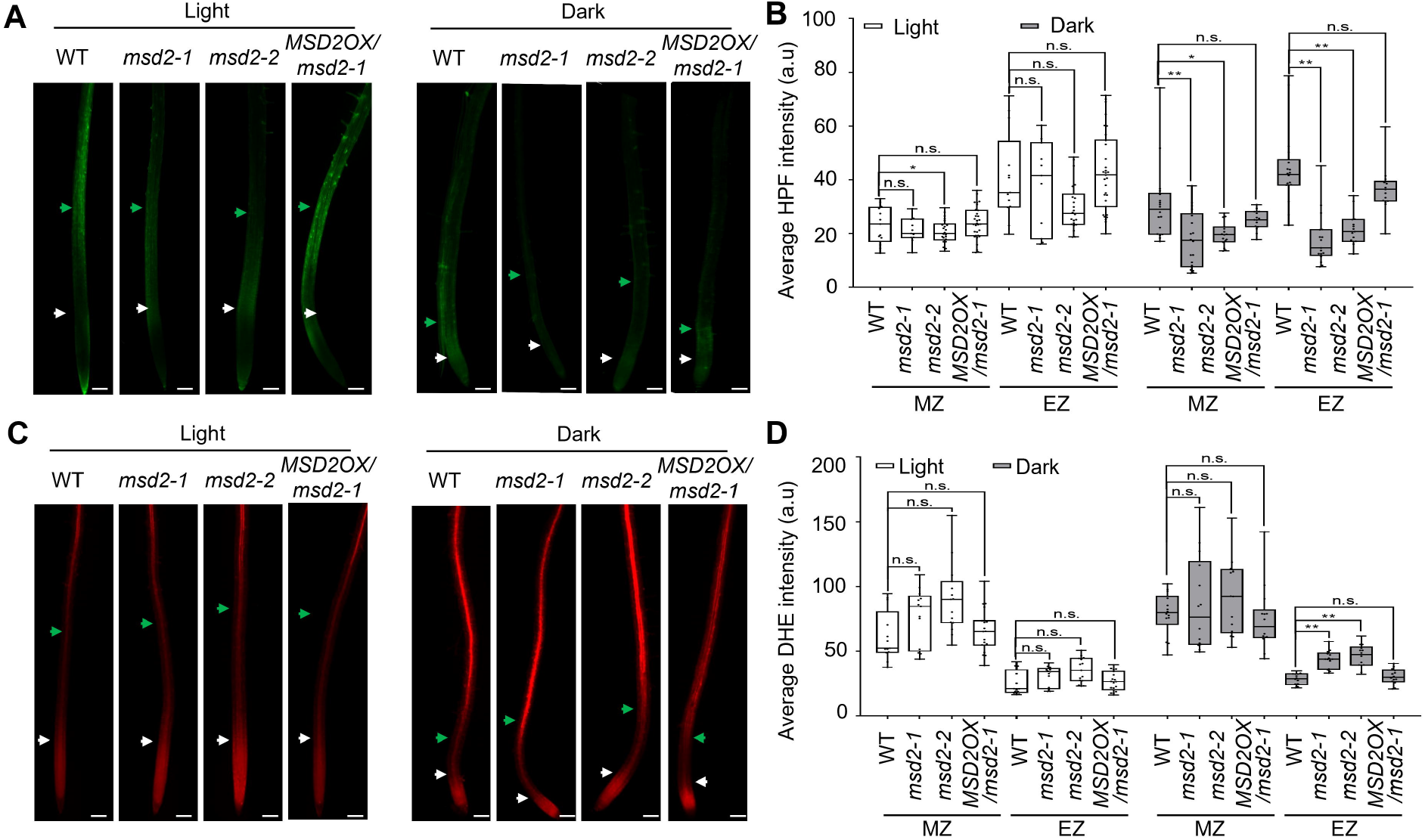
ROS distribution in the WT and *msd2* mutants. (A) Roots of 7-DAG seedlings were stained with hydroxyphenyl fluorescein (HPF). (B) Quantification of HPF fluorescence intensity shown in (A). (C) Roots of 7-DAG seedlings were stained with dihydroethidium (DHE). (D) Quantification of DHE fluorescence intensity shown in (C). *pUBQ10:MSD2-mCherry* was introduced in *msd2-1* mutants (*MSD2OX*/*msd2-1*). Asterisks indicate significant differences (*t*-test, * *P* < 0.05, ** *P* < 0.01). MZ, meristem zone; EZ, elongation zone. Bars = 100 µm.

## 4. Discussion

Among the various ROS-generating machineries, membrane-localized NADPH oxidases have been well studied; key mechanisms regulating their activity and downstream signaling processes are well known [41]. However, how the superoxide generated in the apoplast by NADPH oxidases is delivered into the cell to activate signaling cascades and how specificity is achieved remain unclear. One of the major barriers to our understanding is that the extracellular SOD in Arabidopsis had not been identified. Here, we propose that MSD2 acts as an extracellular SOD, focusing on its role in root morphogenesis.

MSD2 is annotated in TAIR as a member of the Mn-SOD family, but no experimental evidence supports its SOD activity. Here, we experimentally demonstrated that MSD2 has SOD activity and, like MSD1, MSD2 activity was highly resistant to Cu/Zn-SOD or Fe-SOD specific inhibitors (Fig. 2), suggesting that their active centers both incorporate Mn^2+^ in the binding pocket. We supported this hypothesis by comparing the three-dimensional structures of Arabidopsis MSD1 and MSD2: They were highly similar, especially around the Mn^2+^ activity pocket (Fig. 3). The spatial position of the nitrated residue Y63 in MSD1 [18, 42] matched that of Y68 in MSD2 and supported its high nitration probability. MSD2 activity was indeed regulated by nitration (Fig.3 E, F), which provides clues to the extracellular regulatory mechanism of MSD2, although the mechanism of NO synthesis in plants is still not well elucidated.

Notably, MSD1 and MSD2 differed in their subcellular locations. MSD1 localizes to mitochondria [9], whereas MSD2 appeared to be a secreted protein accumulating in the cell wall (Fig. 4). MSD2 might be involved in the metabolism of apoplast ROS under certain conditions. Our promoter analysis in seedlings revealed that *MSD2* is expressed in the junction region of the root and the hypocotyl, the stele of the root mature zone, and the root cap (Fig. 5). These results suggested that *MSD2* expression is highly specific both in space and time and may reflect a critical role at particular growth and developmental stages. Previous studies show the requirement of extracellular SOD in specific cellular processes such as lignin deposition in the Casparian strip formation and floral organ abscission zone development [43, 44]. Peroxidase-dependent lignin polymerization requires extracellular hydrogen peroxide, and cell type–specific lignin deposition is totally abolished in both cell types after treatment with diethyldithiocarbamate (DDC), a Cu/Zn-SOD inhibitor, suggesting critical roles of extracellular SOD in lignin polymerization. However, since DDC does not affect Mn-SOD, it is unlikely that MSD2 contributes to lignin formation. To understand the effects of ROS metabolism mediated by MSD2 on plant development, more in-depth functional studies focusing on the tissues in which *MSD2* is expressed are needed. Analysis of the *MSD2* promoter sequence showed the presence of *cis*-acting elements responding to ABA in addition to those involved in light responses (Fig. 6A). ROS were reported to mediate ABA signaling [45]. Analyzing the effects of ABA on *MSD2* expression and the effect of MSD2 on the ABA signaling pathway would be a good starting point to study the relationship between environmental stress and MSD2-mediated ROS.

During growth, plant development is greatly affected by light, even underground. Light can penetrate several centimeters below the soil surface, and photoreceptors and their associated signal transduction pathways are active in plant roots [46]. Light influences root growth, lateral root formation, and root hair formation, as well as root gravitropic responses, thus guiding the root architecture and growth direction [33]. We discovered that *MSD2* expression is elevated in etiolated seedlings (Fig. 6), suggesting that MSD2 may contribute to ROS metabolism in roots during the light-to-dark transition (Fig. 8). Crosstalk between ROS and light signal pathways was previously reported during leaf development in Arabidopsis [47]. Exogenously applied H_2_O_2_ promoted the establishment of leaves even in the dark, which was accompanied by higher expression of light-responsive genes and a suppression of the inhibitory effects imposed on photomorphogenesis. Little is known about how H_2_O_2_ contributes to root development, which responds differently to light and darkness. As the spatial distribution of O_2_^−^ and H_2_O_2_ strongly influences the spatial transition from proliferation to differentiation in the root [39, 40], increased expression of *MSD2* in the dark may act as an important mediator linking light conditions to root morphogenesis. Phenotypic analysis supported this notion, as the size of the elongation zone was altered in the *msd2* mutant (Fig. 7), along with an altered ROS distribution (Fig. 8). H_2_O_2_ can modulate auxin contents and distribution by affecting the expression of auxin biosynthesis and polar transport genes [48, 49]. It remains to be determined whether and, if so, how altered ROS metabolism by MSD2 might affect auxin flux.

Light activates photoreceptors and their downstream signaling pathways to converge on the master regulators COP1 and HY5. COP1 mediates HY5 degradation in the dark, whereas light blocks the interaction of COP1 with HY5, consequently promoting the accumulation of HY5 and the transcriptional activation of its downstream target genes [36]. Light and dark conditions also alter phytohormone responses, as light facilitates auxin transport from the shoot apex to the root [37], while darkness induces the expression of phytohormone biosynthesis and signaling-related genes for ethylene and gibberellic acid, which regulate skotomorphogenic development [50]. Analysis of *MSD2* expression in a *cop1-4* mutant allele showed that COP1 negatively regulates *MSD2* expression in the dark but had no effect on *MSD2* expression in the light (Fig. 6F). COP1 functions together with SUPPRESSOR OF PHYTOCHROME A-105 1 (SPA1) and related SPA proteins in proteasome-mediated degradation of proteins in response to light exposure. However, a COP1-independent role for SPA proteins was recently suggested [51]. In addition, the rapid light-induced degradation of PHYTOCHROME INTERACTING FACTOR3 (PIF3) was shown to take place in a COP1-independent manner [52]. Future studies investigating the mechanisms regulating the turnover of MSD2 under various light conditions will provide clues to elucidate COP1-independent pathways that have not yet been identified.

## 5. Conclusion

Although it has been suggested that an extracellular SOD may be involved in various signaling pathways [53, 54], its molecular identity had not yet been elucidated in Arabidopsis. We provide experimental evidence that MSD2, which was suggested to be a putative secreted SOD based on sequence homology, possesses SOD activity and is secreted into the vacuole or the extracellular space. Characteristics of *MSD2*, such as its expression in specific tissues or its regulation by light, suggest that it may play an important role in specific signaling processes accompanying ROS metabolism. We propose that the distribution of ROS is in part regulated by MSD2 in roots where it is important to spatially maintain the distribution of different types of ROS [55]. *MSD2* expression is differentially affected by light conditions and consequently contributes to root growth in the dark. These findings not only offer a new component regulating ROS metabolism, but also provide a link between environmental stimuli, ROS metabolism, and plant growth, which can be now further dissected by in-depth studies of the mechanisms of these multifaceted ROS signaling pathways.

## Acknowledgments

We thank June M. Kwak (DGIST) for critical reading of the manuscript and Young Hun Song (Seoul National University) for sharing *cop1-4* mutants. H.C. was supported by the China Scholarship Council (CSC202008140062) and the Natural Science Foundation of China (NSFC31900251). Y.L. was funded by the Suh Kyungbae Foundation (SUHF-19010003) and the National Research Foundation of Korea (NRF-2020R1A2C2013176, and NRF-2021R1A5A1032428). Jinsu L. was funded by the National Research Foundation of Korea (NRF-2020R1I1A1A01068615).

## Author contributions

Y.L. and H.C. conceived the study. H.C., Jinsu L., J.M.L., M.H., Jiyoun L., X.J., and A.E. performed the experiments. H.C. and Y.L. wrote the manuscript. All authors have read and agreed to the published version of the manuscript.

## References

[1] C. Waszczak, M. Carmody, J. Kangasjarvi, Reactive oxygen species in plant signaling, Annu Rev Plant Biol, 69 (2018) 209–236.

[2] K. Mase, H. Tsukagoshi, Reactive Oxygen Species Link Gene Regulatory Networks During Arabidopsis Root Development, Front Plant Sci, 12 (2021) 660274.

[3] R. Mittler, ROS are good, Trends in plant science, 22 (2017) 11–19.

[4] S. Yadav, S.S. Gill, N. Passricha, R. Gill, P. Badhwar, N.A. Anjum, J.-B.J. Francisco, N. Tuteja, Genome-wide analysis and transcriptional expression pattern-assessment of superoxide dismutase (SOD) in rice and Arabidopsis under abiotic stresses, Plant Gene, 17 (2019).

[5] S.S. Gill, N.A. Anjum, R. Gill, S. Yadav, M. Hasanuzzaman, M. Fujita, P. Mishra, S.C. Sabat, N. Tuteja, Superoxide dismutase--mentor of abiotic stress tolerance in crop plants, Environ Sci Pollut Res Int, 22 (2015) 10375–10394.

[6] F. Myouga, C. Hosoda, T. Umezawa, H. Iizumi, T. Kuromori, R. Motohashi, Y. Shono, N. Nagata, M. Ikeuchi, K. Shinozaki, A heterocomplex of iron superoxide dismutases defends chloroplast nucleoids against oxidative stress and is essential for chloroplast development in Arabidopsis, The Plant Cell, 20 (2008) 3148–3162.

[7] M.J. Morgan, M. Lehmann, M. Schwarzlander, C.J. Baxter, A. Sienkiewicz-Porzucek, T.C. Williams, N. Schauer, A.R. Fernie, M.D. Fricker, R.G. Ratcliffe, Decrease in manganese superoxide dismutase leads to reduced root growth and affects tricarboxylic acid cycle flux and mitochondrial redox homeostasis, Plant Physiology, 147 (2008) 101–114.

[8] A.F. Miller, Superoxide dismutases: ancient enzymes and new insights, FEBS Lett, 586 (2012) 585–595.

[9] D.J. Kliebenstein, R.-A. Monde, R.L. Last, Superoxide dismutase in Arabidopsis: an eclectic enzyme family with disparate regulation and protein localization, Plant physiology, 118 (1998) 637–650.

[10] A. Schmidt, M. Gube, A. Schmidt, E. Kothe, In silico analysis of nickel containing superoxide dismutase evolution and regulation, J Basic Microbiol, 49 (2009) 109–118.

[11] P. Dvořák, Y. Krasylenko, M. Ovečka, J. Basheer, V. Zapletalová, J. Šamaj, T. Takáč, In vivo light-sheet microscopy resolves localisation patterns of FSD1, a superoxide dismutase with function in root development and osmoprotection, Plant, Cell & Environment, 44 (2021) 68–87.

[12] C.H. Huang, W.Y. Kuo, C. Weiss, T.L. Jinn, Copper chaperone-dependent and - independent activation of three copper-zinc superoxide dismutase homologs localized in different cellular compartments in Arabidopsis, Plant Physiol, 158 (2012) 737–746.

[13] F. Yamakura, H. Kawasaki, Post-translational modifications of superoxide dismutase, Biochim Biophys Acta, 1804 (2010) 318–325.

[14] J.M. Leitch, C.X. Li, J.A. Baron, L.M. Matthews, X. Cao, P.J. Hart, V.C. Culotta, Post-translational modification of Cu/Zn superoxide dismutase under anaerobic conditions, Biochemistry, 51 (2012) 677–685.

[15] C. Archambaud, M.A. Nahori, J. Pizarro-Cerda, P. Cossart, O. Dussurget, Control of Listeria superoxide dismutase by phosphorylation, J Biol Chem, 281 (2006) 31812–31822.

[16] B. Alvarez, V. Demicheli, R. Durán, M. Trujillo, C. Cerveñansky, B.A. Freeman, R. Radi, Inactivation of human Cu, Zn superoxide dismutase by peroxynitrite and formation of histidinyl radical, Free Radical Biology and Medicine, 37 (2004) 813–822.

[17] V. Demicheli, D.M. Moreno, R. Radi, Human Mn-superoxide dismutase inactivation by peroxynitrite: a paradigm of metal-catalyzed tyrosine nitration in vitro and in vivo, Metallomics, 10 (2018) 679–695.

[18] C. Holzmeister, F. Gaupels, A. Geerlof, H. Sarioglu, M. Sattler, J. Durner, C. Lindermayr, Differential inhibition of Arabidopsis superoxide dismutases by peroxynitrite-mediated tyrosine nitration, J Exp Bot, 66 (2015) 989–999.

[19] M.V. Martin, D.F. Fiol, V. Sundaresan, E.J. Zabaleta, G.C. Pagnussat, oiwa, a female gametophytic mutant impaired in a mitochondrial manganese-superoxide dismutase, reveals crucial roles for reactive oxygen species during embryo sac development and fertilization in Arabidopsis, Plant Cell, 25 (2013) 1573–1591.

[20] T.W. McNellis, A.G. von Arnim, T. Araki, Y. Komeda, S. Miséra, X.W. Deng, Genetic and molecular analysis of an allelic series of cop1 mutants suggests functional roles for the multiple protein domains, The Plant Cell, 6 (1994) 487–500.

[21] M. Martinez-Trujillo, V. Limones-Briones, J.L. Cabrera-Ponce, L. Herrera-Estrella, Improving transformation efficiency of Arabidopsis thaliana by modifying the floral dip method, Plant Mol Biol Rep, 22 (2004) 63–70.

[22] S. Kumar, G. Stecher, M. Li, C. Knyaz, K. Tamura, MEGA X: Molecular Evolutionary Genetics Analysis across Computing Platforms, Mol Biol Evol, 35 (2018) 1547–1549.

[23] K. Arnold, L. Bordoli, J. Kopp, T. Schwede, The SWISS-MODEL workspace: a web-based environment for protein structure homology modelling, Bioinformatics, 22 (2006) 195–201.

[24] D. Wang, D. Liu, J. Yuchi, F. He, Y. Jiang, S. Cai, J. Li, D. Xu, MusiteDeep: a deeplearning based webserver for protein post-translational modification site prediction and visualization, Nucleic Acids Res, 48 (2020) W140–w146.

[25] Z. Liu, J. Cao, Q. Ma, X. Gao, J. Ren, Y. Xue, GPS-YNO2: computational prediction of tyrosine nitration sites in proteins, Mol Biosyst, 7 (2011) 1197–1204.

[26] M. Lescot, P. Déhais, G. Thijs, K. Marchal, Y. Moreau, Y. Van de Peer, P. Rouzé, S. Rombauts, PlantCARE, a database of plant cis-acting regulatory elements and a portal to tools for in silico analysis of promoter sequences, Nucleic acids research, 30 (2002) 325–327.

[27] C. Chen, H. Chen, Y. Zhang, H.R. Thomas, M.H. Frank, Y. He, R. Xia, TBtools: an integrative toolkit developed for interactive analyses of big biological data, Molecular plant, 13 (2020) 1194–1202.

[28] C.A. Schneider, W.S. Rasband, K.W. Eliceiri, NIH Image to ImageJ: 25 years of image analysis, Nat Methods, 9 (2012) 671–675.

[29] W.-Y. Kuo, C.-H. Huang, C. Shih, T.-L. Jinn, Cellular extract preparation for superoxide dismutase (SOD) activity assay, Bio-protocol, 3 (2013) e811.

[30] M. Li, L. Zhu, W. Wang, Improving the thermostability and stress tolerance of an archaeon hyperthermophilic superoxide dismutase by fusion with a unique N-terminal domain, SpringerPlus, 5 (2016) 241.

[31] C.S. Goh, Y. Lee, S.-H. Kim, Calcium could be involved in auxin-regulated maintenance of the quiescent center in the Arabidopsis root, Journal of Plant Biology, 55 (2012) 143–150.

[32] E. Truernit, H. Bauby, K. Belcram, J. Barthélémy, J.C. Palauqui, OCTOPUS, a polarly localised membrane-associated protein, regulates phloem differentiation entry in Arabidopsis thaliana, Development, 139 (2012) 1306–1315.

[33] K. van Gelderen, C. Kang, R. Pierik, Light signaling, root development, and plasticity, Plant physiology, 176 (2018) 1049–1060.

[34] Y. Jing, R. Lin, Transcriptional regulatory network of the light signaling pathways, New Phytol, 227 (2020) 683–697.

[35] R. Podolec, R. Ulm, Photoreceptor-mediated regulation of the COP1/SPA E3 ubiquitin ligase, Curr Opin Plant Biol, 45 (2018) 18–25.

[36] Y. Zhang, C. Wang, H. Xu, X. Shi, W. Zhen, Z. Hu, J. Huang, Y. Zheng, P. Huang, K.X. Zhang, X. Xiao, X. Hao, X. Wang, C. Zhou, G. Wang, C. Li, L. Zheng, HY5 Contributes to Light-Regulated Root System Architecture Under a Root-Covered Culture System, Front Plant Sci, 10 (2019) 1490.

[37] H.-J. Lee, Y.-J. Park, J.-H. Ha, I.T. Baldwin, C.-M. Park, Multiple routes of light signaling during root photomorphogenesis, Trends in plant science, 22 (2017) 803–812.

[38] K. Yokawa, R. Fasano, T. Kagenishi, F. Baluška, Light as stress factor to plant roots– case of root halotropism, Frontiers in plant science, 5 (2014) 718.

[39] A. Eljebbawi, Y. Guerrero, C. Dunand, J.M. Estevez, Highlighting reactive oxygen species as multitaskers in root development, iScience, 24 (2021) 101978.

[40] H. Tsukagoshi, W. Busch, P.N. Benfey, Transcriptional regulation of ROS controls transition from proliferation to differentiation in the root, Cell, 143 (2010) 606–616.

[41] P. Dvorak, Y. Krasylenko, A. Zeiner, J. Samaj, T. Takac, Signaling Toward Reactive Oxygen Species-Scavenging Enzymes in Plants, Front Plant Sci, 11 (2020) 618835.

[42] J. Lozano-Juste, R. Colom-Moreno, J. Leon, In vivo protein tyrosine nitration in Arabidopsis thaliana, J Exp Bot, 62 (2011) 3501–3517.

[43] Y. Lee, M.C. Rubio, J. Alassimone, N. Geldner, A mechanism for localized lignin deposition in the endodermis, Cell, 153 (2013) 402–412.

[44] Y. Lee, T.H. Yoon, J. Lee, S.Y. Jeon, J.H. Lee, M.K. Lee, H. Chen, J. Yun, S.Y. Oh, X. Wen, H.K. Cho, H. Mang, J.M. Kwak, A Lignin Molecular Brace Controls Precision Processing of Cell Walls Critical for Surface Integrity in Arabidopsis, Cell, 173 (2018) 1468–1480 e1469.

[45] J.M. Kwak, I.C. Mori, Z.-M. Pei, N. Leonhardt, M.A. Torres, J.L. Dangl, R.E. Bloom, S. Bodde, J.D.G. Jones, J.I. Schroeder, NADPH oxidase AtrbohD and AtrbohF genes function in ROS-dependent ABA signaling in Arabidopsis, EMBO J, 22 (2003) 2623–2633.

[46] M. Mo, K. Yokawa, Y. Wan, F. Baluska, How and why do root apices sense light under the soil surface?, Front Plant Sci, 6 (2015) 775.

[47] H. Cheng, Q. Liang, X. Chen, Y. Zhang, F. Qiao, D. Guo, Hydrogen peroxide facilitates Arabidopsis seedling establishment by interacting with light signalling pathway in the dark, Plant Cell Environ, 42 (2019) 1302–1317.

[48] A. Huang, Y. Wang, Y. Liu, G. Wang, X. She, Reactive oxygen species regulate auxin levels to mediate adventitious root induction in Arabidopsis hypocotyl cuttings, J Integr Plant Biol, 62 (2020) 912–926.

[49] Y. Fu, Y. Yang, S. Chen, N. Ning, H. Hu, Arabidopsis IAR4 Modulates Primary Root Growth Under Salt Stress Through ROS-Mediated Modulation of Auxin Distribution, Front Plant Sci, 10 (2019) 522.

[50] Deepika, Ankit, S. Sagar, A. Singh, Dark-Induced Hormonal Regulation of Plant Growth and Development, Front Plant Sci, 11 (2020) 581666.

[51] D. Debrieux, M. Trevisan, C. Fankhauser, Conditional involvement of constitutive photomorphogenic1 in the degradation of phytochrome A, Plant Physiol, 161 (2013) 2136–2145.

[52] D. Bauer, A. Viczian, S. Kircher, T. Nobis, R. Nitschke, T. Kunkel, K.C. Panigrahi, E. Adam, E. Fejes, E. Schafer, F. Nagy, Constitutive photomorphogenesis 1 and multiple photoreceptors control degradation of phytochrome interacting factor 3, a transcription factor required for light signaling in Arabidopsis, Plant Cell, 16 (2004) 1433–1445.

[53] O. Pechanova, C.-Y. Hsu, J.P. Adams, T. Pechan, L. Vandervelde, J. Drnevich, S. Jawdy, A. Adeli, J.C. Suttle, A.M. Lawrence, T.J. Tschaplinski, A. Séguin, C. Yuceer, Apoplast proteome reveals that extracellular matrix contributes to multistress response in poplar, BMC Genomics, 11 (2010) 674.

[54] F.A. Kaffarnik, A.M. Jones, J.P. Rathjen, S.C. Peck, Effector proteins of the bacterial pathogen Pseudomonas syringae alter the extracellular proteome of the host plant, Arabidopsis thaliana, Mol Cell Proteomics, 8 (2009) 145–156.

[55] X. Zhou, Y. Xiang, C. Li, G. Yu, Modulatory Role of Reactive Oxygen Species in Root Development in Model Plant of Arabidopsis thaliana, Front Plant Sci, 11 (2020) 485932.

[56] J.J.A. Armenteros, K.D. Tsirigos, C.K. Sønderby, T.N. Petersen, O. Winther, S. Brunak, G. von Heijne, H. Nielsen, SignalP 5.0 improves signal peptide predictions using deep neural networks, Nature biotechnology, 37 (2019) 420–423.

