## Supplemental Information for "MSD2, an apoplastic Mn-SOD, contributes to root skotomorphogenic growth by modulating ROS distribution in Arabidopsis"

**Supplementary data**

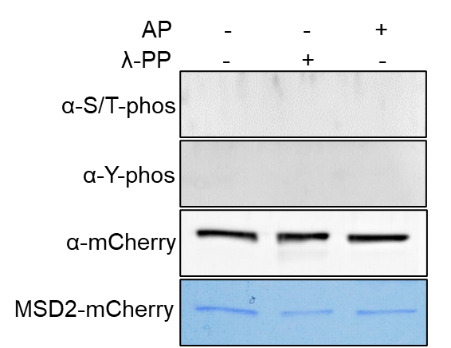

**Fig. S1.** Assessment of MSD2 phosphorylation after pull-down by immunoblotting with anti-Ser/Thr and anti-Tyr phosphorylation-specific antibodies. Total proteins were extracted from transgenic plants expressing *pUBQ10:MSD2-mCherry*. AP, alkaline phosphatase; λ-PP, lambda protein phosphatase.

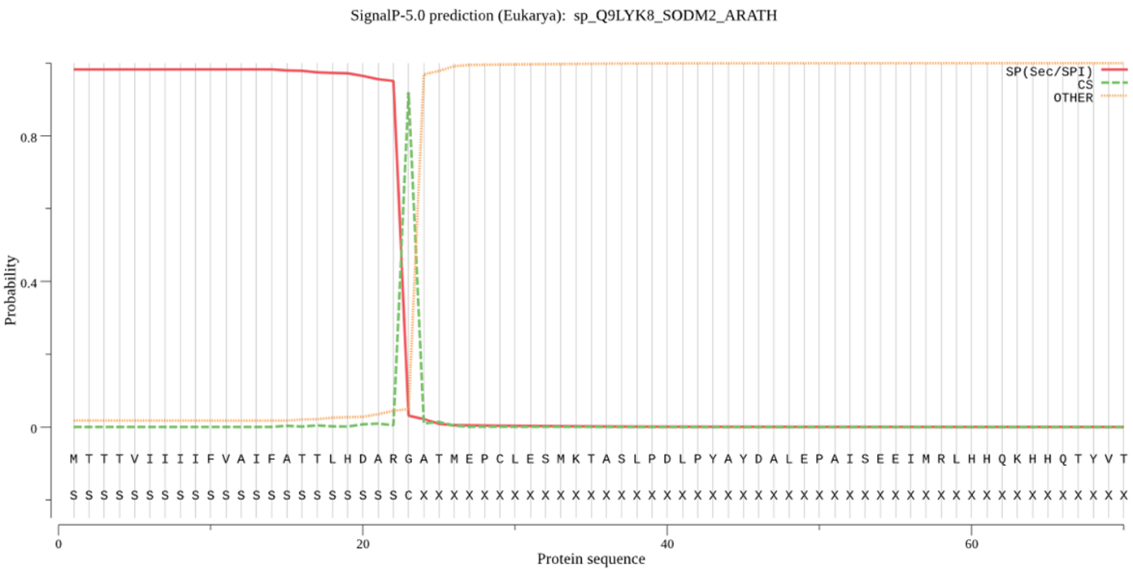

**Fig.S2.** *In silico* prediction of a signal peptide in the N terminus of MSD2. The red line indicates the possible signal peptide, according to SignalP-5.0 [55].

**Table S1.** List of SODs used for phylogenetic tree analysis.

| **Locus** | **Definition** | **Sequence** |
| --- | --- | --- |
| NP_001077494.1 | CSD 1 Arabidopsis thaliana | MAKGVAVLNSSEGVTGTIFFTQEGDGVTTVSGTVSGLKPGLHGFHVHALGDTTNGCMSTGPHFNPDGKTHGAPEDANRHAGDLGNITVGDDGTATFTITDCQIPLTGPNSIVGRAVVVHADPDDLGKGGHELSLATGNAGGRVACGIIGLQG |
| AZP89710.1 | SOD Caragana korshinskii | MAKGVAVLNSSEGVTGTIFFTQEGDGVTTVSGIVSGLKPGLHGFHVHALGDTTNGCMSTGPHFNPDGKTHGAPEDANRHAGDLGNITVGDDGTATFTITDCQIPLTGPNSIVGRAVVVHADPDDLGKGGHELSLATGNAGGRVACGIIGLQG |
| XP_002889716.1 | CSD1 Arabidopsis lyrata subsp. lyrata | MAKGVAVLNSSEGVKGTIFFTQEGDGVTTVTGTVSGLKPGLHGFHVHALGDTTNGCMSTGPHFNPDGKTHGAPEDANRHAGDLGNITVGDDGTATFTITDTQIPLTGPNSIVGRAVVVHADPDDLGKGGHELSLATGNAGGRVACGIIGLQG |
| KAG7653595.1 | CSD Arabidopsis suecica | MAKGVAVLNSSEGVKGTIFFTQEGDGVTTVTGTVSGLKPGLHGFHVHALGDTTNGCMSTGPHFNPDGKTHGAPEDANRHAGDLGNITVGDDGTASFTITDTQIPLTGPNSIVGRAVVVHADPDDLGKGGHELSLATGNAGGRVACGIIGLQG |
| XP_013638847.1 | CSD1 Brassica oleracea var. oleracea | MGKGVAVLNSSEGVKGTIFFTQEGDGVTSVTGTVSGLKPGLHGFHVHALGDTTNGCMSTGPHFNPDGKQHGAPEDANRHAGDLGNIIVGDDGTATFTITDCQIPLSGPNSIVGRAVVVHADPDDLGKGGHELSLATGNAGGRVACGIIGLQG |
| XP_006305743.1 | CSD1 Capsella rubella | MAKGVAVLNSSEGVKGTIFFTQEGQGVTTVTGTLSGLKPGLHGFHVHALGDTTNGCMSTGPHFNPDGKTHGAPEDATRHAGDLGNITVGDDGTATFTITDCQIPLTGPNSIVGRAVVVHADPDDLGKGGHELSLTTGNAGGRIACGIIGLQG |
| XP_009148138.1 | CSD1 Brassica rapa | MGKGVAVLNSGEGVKGTIFFTQEGDGVTTVTGTVSGLKPGLHGFHVHALGDTTNGCMSTGPHFNPDGKQHGAPEDANRHAGDLGNIIVGDDGTATFTITDCQIPLSGPNSIVGRAVVVHADPDDLGKGGHELSLATGNAGGRVACGIIGLQG |
| XP_006417641.1 | CSD1 Eutrema salsugineum | MAKGVAVLSSSEGVKGTIFFTQEGQGETTVSGTVSGLKPGLHGFHVHALGDTTNGCMSTGPHFNPDGKQHGAPEDANRHAGDLGNIVVGDDGTATFSITDCQIPLTGPNSIIGRAVVVHADPDDLGKGGHELSLATGNAGGRVACGIIGLQG |
| XP_018486007.1 | CSD1 Raphanus sativus | MGKGVAVLNSSEGVKGTIFFTQEGNGATTVTGTVSGLKPGLHGFHVHALGDTTNGCMSTGPHFNPDGKTHGAPEDANRHAGDLGNITVGDDGTATFTITDSQIPLDGPNSIVGRAVVVHADPDDLGKGGHELSLATGNAGGRVACGIIGLQG |
| AAD05576.1 | CSD Raphanus sativus | MGKGVAVLNSSEGVKGTIFFTQEGNGSTTVTGTVSGLKPGLHGFHVHALGDTTNGCMSTGPHFNPDGKTHGAPEDANRHAGDLGNITVGDDGTASFTITDSQIPLDGPNSIVGRAVVVHADPDDLGKGGHELSLATGNAGGRVACGIIGLQG |
| XP_013604816.1 | CSD Brassica oleracea var. oleracea | MAKGVAVLNSSEGVKGTIFFTHEGNGATTVTGTVSGLKPGLHGFHVHALGDTTNGCMSTGPHFNPDGKTHGAPEDANRHAGDLGNIIVGDDGTATFTITDSQIPLSGPNSIVGRAVVVHADPDDLGKGGHELSLSTGNAGGRVACGIIGLQG |
| AYF57908.1 | CSD2 Brassica oleracea var. botrytis | MAKGVAVLNSSEGVKGTIFFTHEGNGATTVTGTVSGLRPGLHGFHVHALGDTTNGCMSTGPHFNPDGKTHGAPEDANRHAGDLGNIIVGDDGTATFTITDSQIPLSGPNSIVGRAVVVHADPDDLGKGGHELSLSTGNAGGRVACGIIGLQG |
| AAC25568.1 | CSD Brassica rapa subsp. pekinensis | MAKGVAVLNSSEGVKGTIFFTQEGDGATTVTGTVSGLKPGPHGFHVHALGDTTNGCMSTGPHFNPDGKTHGAPEDANRHAGDLGNIIVGDDGTATFTITDSQIPLTGPNSIVGRAVVVHADRDDLGKGGHELSLSTGNAGGRVACGIIGLQG |
| XP_009118346.1 | CSD Brassica rapa | MAKGVAVLNSSEGVKGTIFFTQEGDGATTVTGTVSGLKPGPHGFHVHALGDTTNGCMSTGPHFNPDGKTHGAPEDANRHAGDLGNIIVGDDGTATFTITDSQIPLTGPNSIVGRAVVVHAERDDLGKGGHELSLSTGNAGGRVACGIIGLQG |
| P09678.2 | CSD Brassica oleracea var. capitata | MAKGVAVLNSSEGVKGTIFFTHEGNGATTVTGTVSGLRPGLHGFHVHALGDNTNGCMSTGPHFNPDGKTHGAPEDANRHADLGNIIVGDDGTATFTITDSQIPLSGPNSIVGRAIVVHADPDDLGKGGHELSLSTGNAGGRVACGIIGLQG |
| Q42612.3 | CSD2 Brassica juncea | MGKGVAVLNSSEGVKGTIFFAQEGEGKTTVTGTVSGLKPGLHGFHVHALGDTTNGSMSTGPHFNPDGKQHGAPEDANRHAGDLGNIIVGDDGTATFTITDCQIPLSGPNSIVGRAVVVHADPDVLGKGGHELSLTTGNAGGRVACGIIGLQG |
| AKN08991.1 | SOD Luffa aegyptiaca | MVKGVAVLGSSEGVSGTIFFSQEGDGPTTVTGNVSGLKPGLHGFHVHALGDTTNGCMSTGPHFNPAGKQHGAPEDENRHAGDLGNITVGEDGKASFTISDCQIPLSGPNSIIGRAVVVHGDPDDLGKGGHELSLTTGNAGGRVACGIIGLQG |
| XP_038990856.1 | CSD Hibiscus syriacus | MVKAVAVLSSSEGVKGTVFFTQEGDGPTTVTGNISGLKPGLHGFHVHALGDTTNGCMSTGPHFNPAGKEHGAPEDENRHAGDLGNVTVGDDGTASFSITDNQIPLTGPNSIIGRAVVVHADPDDLGKGGHELSKSTGNAGGRVACGIIGLQG |
| XP_019440527.1 | CSD Lupinus angustifolius | MAKGVAVLGSSEGVAGTVYFIQEGDGPTTVTGTLSGLKPGLHGFHVHALGDTTNGCLSTGPHYNPNGKEHGAPEDDNRHAGDLGNINVGDDGTVTFSITDSQIPLTGPNSIIGRAVVVHADPDDLGKGGHELSKATGNAGGRVACGIIGLQG |
| XP_030467015.1 | CSD 1 Syzygium oleosum | MVKAVAVLGSSEGVTGTIYFTQEGDAPTTVTGSLSGLKPGIHGFHVHALGDTTNGCMSTGPHFNPAGKEHGAPEDENRHAGDLGNVTVGDDGTATFTIIDKQIPLTGPNSIVGRAVVVHADPDDLGKGGHELSKTTGNAGGRVACGIIGLQG |
| XP_006382702.2 | CSD 2 Populus trichocarpa | MVKAVAVLNSSEGVSGTIFFTQEGDGQTTVTGNLSGLKPGLHGFHVHALGDTTNGCMSTGPHFNPVGKEHGAPEDENRHAGDLGNVTVGDDGTATFTIIDKQIPLTGPHSIIGRAVVVHGDPDDLGKGGHELSKTTGNAGGRVACGIIGLQG |
| ACI46676.1 | CSD Gossypium arboreum | MVKAVAVLSSSEGVSGTVFFTQEGDGPTTVTGNLSGLKPGLHGFHVHALGDTTNGCMSTGPHFNPAGKEHGAPEDVNRHAGDLGNVTVGDDGCASFSITDKQIPLTGPNSIIGRAVVVHADPDDLGKGGHELSKSTGNAGGRVACGIIGLQG |
| NP_565666.1 | CSD2 Arabidopsis thaliana | MAATNTILAFSSPSRLLIPPSSNPSTLRSSFRGVSLNNNNLHRLQSVSFAVKAPSKALTVVSAAKKAVAVLKGTSDVEGVVTLTQDDSGPTTVNVRITGLTPGPHGFHLHEFGDTTNGCISTGPHFNPNNMTHGAPEDECRHAGDLGNINANADGVAETTIVDNQIPLTGPNSVVGRAFVVHELKDDLGKGGHELSLTTGNAGGRLACGVGLTPL |
| NP_197311.1 | CSD3 Arabidopsis thaliana | MEAPRGNLRAVALIAGDNNVRGCLQFVQDISGTTHVTGKISGLSPGFHGFHIHSFGDTTNGCISTGPHFNPLNRVHGPPNEEERHAGDLGNILAGSNGVAEILIKDKHIPLSGQYSILGRAVVVHADPDDLGKGGHKLSKSTGNAGSRVGCGIIGLQSSADAKL |
| XP_034917943.1 | CSD2 Populus alba | MVKAVAVLNSSEGVSGTIFFTQEGDGPTTVTGNLSGLKPGLHGFHVHALGDTTNGCMSTGPHFNPVGKEHGAPEDENRHAGDLGNVTVGDDGTATFTIIDKQIPLTGPHSIIGRAVVVHGDPDDLGKGGHELSKTTGNAGGRVACGIIGLQG |
| XP_010025114.2 | CSD Eucalyptus grandis | MVKAVAVLGSSEGVTGTVHFVQEGDGPTTVTGSLSGLKPGLHGFHVHALGDTTNGCMSTGPHFNPSGKEHGAPEDENRHAGDLGNVTVGDNGTATFTITDKQIPLTGPNSIVGRAVVVHSDPDDLGKGGHELSKTTGNAGGRVACGIIGLQG |
| AAD01605.1 | CSD Populus tremuloides | MVKAVAVLNSSEGVSGTIFFTQEGDGPTTVTGNLSGLKPGLHGFHVHALGDTTNGCMSTGPHFNPVGKEHGAPEDENRHAGDLGNVTVGDDGTAAFTIIDFQIPLTGPHSIIGRAVVVHGDPDDLGKGGHELSKTTGNAGGRVACGIIGLQG |
| Q42611.3 | CSD 1 Brassica juncea | MGKGVRVLNSSEGVKGTIFFTQEGNGTTTVTGTVSGLKPGLHGFHVHALGDTTNGCMSTGPHFNPEGKTHGAPEDANRHAGDLGNITVGDDGTATFTITDSQIPLDGPNSIVGRAVVVHAEPDDLGKGGHELSLTTGNAGGRVACGIIGLQG |
| XP_024186633.1 | CSD Rosa chinensis | MAKGVAVLCSSEGVKGTILFTQEGDGPTTVTGNVSGLKPGLHGFHVHALGDTTNGCMSTGPHFNPAAKEHGAPEDENRHAGDLGNITVGDDGTATFTIVDKQIPLTGPHSIIGRAVVVHGDPDDLGKGGHELSKSTGNAGGRIACGIIGLQG |
| BAJ07302.1 | CSD Melastoma malabathricum | MVKAVVVLGNSEGVSGTVYFTQEGDGPTTVTGSLSGLKPGLHGFHVHALGDTTNGCMSTGPHFNPAGKEHGAPEDENRHAGDLGNVTVGDDGTATFTITDKQIPLFGPNSIIGRAVVVHADPDDLGKGGHELSKSTGNAGGRIACGIIGLQG |
| ABF48717.1 | CSD Populus suaveolens | MVKAVAVLNSSEGVSGTIFFTQEGDGPTTVTGNLSGLKPGLHGFHVHALRDTTNGCMSTGPHFNPVGKEHGAPEDENRHAGDLGNVTVGDDGTATFTIIDKQIPLTGPHSIIGRAVVVHGDPDDLGKGGHELSKTTGNAGGRVACGIIGLQG |
| XP_022158885.1 | CSD Momordica charantia | MVKAVAVLGASDVVSGTIFFSQEGDGPTTVTGNVSGLKAGLHGFHVHALGDTTNGCMSTGPHFNPAGKQHGAPEDENRHAGDLGNITVGEDGKASFTITDCQIPLSGPNSIIGRAVVVHADPDDLGKGGHELSLTTGNAGGRVACGIIGLQG |
| ACB38158.1 | CSD Potentilla atrosanguinea | MAKGVAVLSSSEGVAGTILFTQEGDGPTTVTGNISGLKPGLHGFHVHALGDTTNGCMSTGPHFNPAGKEHGSPEDETRHAGDLGNITVGDDGTACFTIVDKQIPLTGPHSIIGRAVVVHADPDDLGKGGHELSKSTGNAGGRIACGIIGLQG |
| XP_010674951.1 | CSD Beta vulgaris subsp. vulgaris | MGKAVVVLNSSEGVTGTIYFTQEGDGPTTVTGNISGLKPGLHGFHVHALGDTTNGCMSTGPHYNPAGKEHGAPEDEVRHAGDLGNVTVGDDGTATFTIIDSQIPLCGPNSVVGRAVVVHADPDDLGRGGHELSKTTGNAGGRVACGVIGLQG |
| XP_021773039.1 | CSD Chenopodium quinoa | MGKAVVVLSSSEGVAGTVYFTQEGDGPTTVTGNVSGLKPGLHGFHVHALGDTTNGCMSTGPHFNPNGKEHGAPEDEVRHAGDLGNITVGDDGTATFTIIDNQIPLSGPNSIVGRAVVVHADPDDLGRGGHELSKTTGNAGGRIACGIIGLQG |
| XP_030534739.1 | CSD Rhodamnia argentea | MVKAVAVLGSSEGVSGTIYFTQEGDAPTTVTGSLSGLKPGLHGFHVHALGDTTNGCMSTGPHFNPAGKEHGAPEDENRHAGDLGNVTVGDDGAATFTIIDKQIPLAGPNSIVGRAVVVHTDPDDLGKGGHELSKTTGNAGGRVACGIIGLQG |
| APT43083.1 | CSD Solanum melongena | MVKAVAVLNSSEGVSGTIFFTQEADGPTTVSGNISGLKPGLHGFHVHALGDTTNGCMSTGPHFNPAGKDHGAPEDELRHAGDLGNITAGEDGTASFTITDKQIPLTGSQSIIGRAVVVHADPDDLGKGGHELSKSTGNAGGRIACGIIGLQG |
| XP_011032547.1 | CSD 2 Populus euphratica | MVKAVAVLNSSEGVNGTIFFTQEGDGPTTVTGNLSGLKPGPHGFHVHALGDTTNGCMSTGPHFNPLGKEHGAPEDENRHAGDLGNVTVGDDGTATFTIIDKQIPLTGPHSIIGRAVVVHGDPDDLGKGGHELSKTTGNAGGRVACGIIGLQG |
| AFF57842.1 | CSD Tetradium ruticarpum | MVKAVAVLGSSEGVKGTVSFTQEGDGPTTVTGSLSGLKPGLHGFHVHALGDTTNGCMSTGPHFNPAGKEHGAPEDENRHAGDLGNVNVGDDGTATFTIVDNQIPLSGPNSIIGRAVVVHADPDDLGKGGHELSKTTGNAGGRVACGIIGLQG |
| XP_034920401.1 | CSD Populus alba | MVKAVAVLNSSEGVKGTINFTQEGDGPTTVTGSLCGLKPGLHGFHVHALGDTTNGCMSTGPHFNPVGKEHGAPEDENRHAGDLGNVTVGDDGTATVSIIDNQIPLTGPNSIVGRAVVVHADPDDLGKGGHELSKSTGNAGGRVACGVIGLQG |
| XP_009624899.1 | CSD Nicotiana tomentosiformis | MVKAVAVLSSSEGVSGTIFFTQDGDAPTTVTGNVSGLKPGLHGFHVHALGDTTNGCMSTGPHYNPAGKEHGAPEDEVRHAGDLGNITVGEDGTASFTITDKQIPLAGPQSIIGRAVVVHADPDDLGKGGHELSKATGNAGGRVACGIIGLQG |
| XP_012464569.1 | CSD Gossypium raimondii | MVKAVAVLSSNEGVSGTVFFSQEGDGPTTVTGNLSGLKAGLHGFHVHALGDTTNGCMSTGPHFNPAGKEHGAPEDENRHAGDLGNVTVGDDGCASFSITDKQIPLTGPNSIIGRAVVVHADPDDLGKGGHELSKSTGNAGGRVACGIIGLQG |
| XP_020873684.1 | FSD1 Arabidopsis lyrata subsp. lyrata | MYDATRLPKSRAIHKSQNTDSTGVCSPHLSLKLELQFKRMAASSAVTANYVLKPPPYALDALEPHMSKQTLEFHWGKHHRAYVENLKKQVLGTELEGKPLEHIIHSTYNNGELLPAFNNAAQAWNHEFFWESMKPGGGGKPSGELLALLERDFTSYEKFYEEFNAAAATQFGAGWAWLAYANDKLKVVKTPNAVNPLVLGSFPLLTIDVWEHAYYLDFQNRRPDYIKTFMTNLVSWEAVSARLEAAKAASSSSS |
| XP_010448478.1 | FSD1 Camelina sativa | MHLNYQKAEPSIKLKTHTDSTGVCSPHLSLKLELQSQSMAASIAVTANYVLQPPPYALDALEPHMSKQTLEFHWGKHHRAYVDNLKKQVLGTDLEGKPLDHIINKTYNNGDLLPAFNNAAQAWNHEFFWESMKPNGGGKPSGELLALLERDFTSYEKFYEEFNAAAATQFGAGWAWLAYANDKLKVVKTPNAVNPLVLGSFPLLTIDVWEHAYYLDFQNRRPDYIKTFMNNLVSWEAVSARLEAAKVASSSSA |
| XP_013608657.1 | FSD1 Brassica oleracea var. oleracea | MAASAAVTANYVLKPPPYPLDALEPHMSKQTLEFHWGKHHRAYVDNLKKQVLGSELEGKPLEHIIQNTYNNGDLLPPFNNAAQAWNHEFFWESMKPGGGGKPSGELLALLERDFTSYEKFYDEFNAAAATQFGAGWAWLAYADNKLKVVKTPNAVNPLVLGSFPLLTIDVWEHAYYLDFQNRRPDYIKTFMNNLVSWEAVSSRLEAAKAASS |
| XP_018472329.1 | FSD1 Raphanus sativus | MAASAAVTANYVLKPPPYPLDALEPHMSKQTLEFHWGKHHRAYVDNLKKQVLGSELEGKPLEHIIQNSYNNGDLLPPFNNAAQAWNHEFFWESMKPGGGGKPSGELLALLERDFTSYEKFYDEFNAAAATQFGAGWAWLAYADNKLKVVKTPNAVNPLVLGSFPLLTIDVWEHAYYLDFQNRRPDYIKTFMNNLVSWEAVSSRLEAAKAASA |
| AEG89520.1 | FSD1 Brassica oleracea var. botrytis | MAASAAVTANYVLKPPPYPLDALEPHMSKQTLEFHWGKHHRAYVDNLKKQVLGSELEGKPLEHIIQNTYNNGDLLPAFNNAAQAWNHEFFWESMKPGGGGKPSGELLALLERDFTSYEKFYDEFNAAAATQFGAGWAWLAYADNKLKVMKTPNAVNPLVLGSFPLLTIDFWEHAYYLDFQNRRPDYIKTFMNNLVSWEAVSSRLEAAKAASS |
| XP_006282692.2 | FSD1 Capsella rubella | MAASTAVTANYVLKPPPYALDALEPHMSKQTLEFHWGKHHRAYVENLKKQVLETELEGKPLDHIIHKTYNNGDLLPAFNNAAQAWNHEFFWESMKPGGGGKPSGELLALLERDFTSYEKFYEEFNAAAATQFGAGWAWLAYANDKLKVVKTPNAVNPLVLGSFPLLTIDVWEHAYYLDFQNRRPDYIKTFMNNLVSWEAVSARLEAAKAASSSSA |
| XP_006413341.1 | FSD1 Eutrema salsugineum | MAASNTVTANYVLKPPPYSLDALEPHMSRQTLEFHWGKHHRAYVDNLKKQVLGTELEGKPLDHIILKTYNNGDLLPAFNNAAQAWNHEFFWESMKPGGGGKPSGELLALLERDFTSYEKFYDEFNAAAATQFGAGWAWLAYANDKLKVVKTPNAVNPLVLDSFPLLTIDVWEHAYYLDFQNRRPDYIKTFMNNLVSWEAVSSRLEAAKAASASSA |
| OAO98056.1 | FSD1 Arabidopsis thaliana | MAASSAVTANYVLKPPPFALDALEPHMSKQTLEFHWGKHHRAYVDNLKKQVLGTELEGKPLEHIIHSTYNNGDLLPAFNNAAQAWNHEFFWESMKPGGGGKPSGELLALLERDFTSYEKFYEEFNAAAATQFGAGWAWLAYSNEKLKVVKTPNAVNPLVLGSFVSFSLRNLNTQFT |
| XP_021617104.1 | FSD Manihot esculenta | MATTAATATSLNCARFPRQAGRLNQATRGFQWTKEIHCTTKARPAIITAKFELKPPPYPLNALEPHMSKDTLEFHWGKHHRAYVDNLNKQIVGTELDSKPLEDVVIATYNKGDVLPAFNNAAQAWNHEFFWGCMKPCGGGKPSGELLQLIERDFGSFEKFVEEFKSAAATQFGSGWAWLVYKTDKLDVENAVNPRPSEEDKKLAVLKSPNAVNPLVWDYSPLLTIDVWEHAYYLDFQNRRPDYISTFLEKLVSWEAVSCRLEAAKAQAAGNS |
| XP_015079796.1 | FSD Solanum pennellii | MAATPSANSLTSAFLPPQGLNGSSKSLQWRTQKLQKQFGRKAGSATITAKFDLNPPPYPMDALEPHMSNKTFEFHWGKHHRAYVDNLNKQIDGTELDGKTLEDIILVTYNNGAPLPAFNNAAQAWNHQFFWESMKPNGGGEPSGELLELINRDFGSYDTFVKEFKAAAATQFGSGWAWLAYKPEDKKLALVKTPNAENPLVLGYTPLLTIDVWEHAYYLDFQNRRPDYISIFMEKLVSWEAVSCRLKAATA |
| XP_031277304.1 | FSD Pistacia vera | MASLATRPIAFTFPCQGLGGCTPGLQFQWTNKKMVPRKCARKAGSTKICAKFELKPPPYPMDALEPHMSRETLEYHWGKHHRAYVENLNKQIVGTELDGMSLGDIVIVSYNNGDMLPAFNNAAQAWNHEFFWESMKPSGGGKPSGELLQLIERDFGSFEKFLKEFKSAAATQFGSGWAWLAYKANRLNVDNAVNPLPSEEDKKLVVVKSPNAVNPLVWDYFPLLTIDVWEHAYYLDFQNRRPDYISIFMDKLVSWEAVSRRLEIAKALLVEREKEEARKEREEEDEEMADDEAVKMYVESDGDDPEND |
| XP_019246046.1 | FSD Nicotiana attenuata | MAATTASANSLTSAFLPRLGFHGSSHQSLQLRSQKFARKAGSGTITAKFELQPPPYPMDALEPHMSSRTFEFHWGKHHRAYVDNLNKQIDGTELDGKTLEDIILVTYNKGAPLPAFNNAAQAWNHQFFWESMKPNGGGEPSGELLELINRDFGSYDAFVKEFKAAAATQFGSGWAWLAYKPEEKKLALVKTPNAENPLVLGYTPLLTIDVWEHAYYLDFQNRRPDYISIFMEKLVSWEAVSSRLKAATA |
| AAQ18699.1 | FSD Solanum lycopersicum | MAATASANSLTSAFLPPQGFNGSSKSLQWRTQKKQFGRKAGSATITAKFDLIPPPYPMDALEPHMSSRTFEFHWGKYHRAYVDNLNKQIDGTELDGKTLEDIILVTYNNGAPLPAFNNAAQAWNHQFFWESMKPNGGGEPSGELLELINRDFGSYDTFVKEFKAAAATQFGSGWAWLAYKPEDKKLALVKTPNAENPLVLGYTPLLTIDVWEHAYYLDFQNRRPDYISIFMEKLVSWEAVSIRLKAASA |
| XP_009586624.1 | FSD Nicotiana tomentosiformis | MMMAATTASANSLTSAFLPRLGFHGSSHKSLQLRTQKKQFARKAGSGTITAKFELQSPPYPMDALEPHMSSRTFEFHWGKHHRAYVDNLNKQIDGTELDGKTLEDIILVTYNKGAPLPAFNNAAQAWNHQFFWESMKPNGGGEPSGELLELINRDFGSYDAFVKEFKAAAATQFGSGWAWLAYKPEEKKLALVKTPNAENPLVLGYTPLLTIDVWEHAYYLDFQNRRPDYISIFMEKLVSWEAVSSRLKAATA |
| PHT44981.1 | FSD2 Capsicum baccatum | MAAANSLSLTSAFLPRHGFHGPSHNQNVQLSTQKKQFARKAGSGTITAKFELQPPPYPMDALEPHMSSRTFEFHWGKHHRAYVDNLNKQIDGTELDGKTLEDIILITYNKGSPLPAFNNAAQAWNHQFFWESMKPNGGGEPSGELLELINRDFGSYDTFVKEFKAAAATQFGSGWAWLAYKPEDKKLTLVKTPNAENPLVLGYTALLTIDVWEHAYYLDFQNRRPDYISIFMEKLVSWEAVSSRLKTATA |
| XP_016575659.1 | FSD Capsicum annuum | MAAANSLSLTSAFLPRHGFHGPSHNQNVQLSTQKKQFARKAVSGTITAKFELQPPPYPMDALEPHMSSRTFEFHWGKHHRAYVDNLNKQIDGTELDGKTLEDIILITYNKGSPLPAFNNAAQAWNHQFFWESMKPNGGGEPSGKLLELINRDFGSYDTFVKEFKAAAATQFGSGWAWLAYKPEDKKLALVKTPNAENPLVLGYTALLTIDVWEHAYYLDFQNRRPDYISIFMEKLVSWEAVSSRLKTATA |
| KAF3782520.1 | FSD Nymphaea thermarum | MESAALSALPSQMLPLSCSGLKKSPFAFQLKLSAPKGTRLRKRCAPITAQFELKPPPYSLDALEPHMSRNTFEFHWGKHHRAYVDNLNKQIQGTELDGKSLEEIIVITYNKGDPLPPFNNAAQVWNHDFFWECMKPGGGGVPSGTLLELIKRDFGSYDAFLKEIKAAAATQFGSGWAWLTYNGDKLEVPKLGIAKTPNAVNPLVWDNTTPLLTIDVWEHAYYLDYQNRRPDYVSIFLENLVSWEAVSARLEAAKLKVAA |
| PON71074.1 | M/FSD Parasponia andersonii | MVALAPLASTTSTTKPYSLTCALFPSRQGLSGNFGSSRTRRLKMSSDASKAVAKIELRPPPYPLHGLEPHMSKNTLEFHWGKHHRAYVDNLNKQIVGTELDGLPLEEIIVRTYNKGDLLPPFNNAAQIWNHDFFWESMKPGGGGKPSGELLRLIERDFGSFDSFIGEFKTAAATQFGSGWAWLAFKDSKLIIVKTPNAVNPLVLDSYVSFPLLTIDVWEHAYYLDFQNRRPDYISLFVEKLVSWDSVSARLEAAKAQAA |
| PON92464.1 | MSD Trema orientale | MVAVAPVASTTSTSKPYSLTCALFPSRQGLSGNFGSFRTRRLKMSSGASKAVAKIELRPPPYPLHGLEPHMSKNTLEFHWGKHHRAYVDNLNKQIVGTELDGLPLEEIIVRTYNKGDLLPPFNNAAQIWNHDFFWESMKPGGGGKPSGELLQLIERDFGSFDSFIGEFKTAAATQFGSGWAWLAFKDSKLIIVKTPNAVNPLVLDSYVSFPLLTIDVWEHAYYLDFQNRRPDYISLFVEKLVSWDSVNARLEAAKAQAA |
| OMO52322.1 | MSD Corchorus olitorius | MAAAASSSMATSRRFSRKAGSNLITANFELKPPPYPLNALEPHMSRETLEYHWGKHHRAYVDNLNKQIVGTGLEGLSLEDTIIVTYNKGDMLPAFNNAAQAWNHEFFWESIKPGGGGKPSGELLDLIERDFGSFEQFIQEFKSAAATQFGSGWAWLAYKANRLDMENAVNPWPSDKDKKLVIVKSPNAVNPLVWDYFPLLTIDVWEHAYYLDFQNRRPDYISMFMEKLVSWEAVSARLEKAKAQAAEREIEEERRRKEEEEEQSDDDDVQLYLDSDTDDSDSE |
| XP_017972312.1 | FSD Theobroma cacao | MAAAASMATSLTFPLLPSQAGLRGPFTSSLPCTIPQRRFWRKVATNLITAKFELKPPPYTLNALEPHMSRQTLEYHWGKHHRTYVENLNKQIAGTELEGLPLEDIIIVSYNNGDILPAFNNAAQAWNHDFFWESMKPGGGGKPSGDLLDLIERDFGSFEQFIQEFKSAAAAQFGSGWAWLAYKANRLDVENAVNPWPSEKDKKLVVVKSPNAVNPLVWDYFPLLTIDVWEHAYYLDFQNRRPDYISMFMEKLISWEAVSARLEKAKALAAEREMEEERRKKEEEEKQTDDEAVEMYLDSDTDDSEAE |
| NP_199923.1 | FSD2 Arabidopsis thaliana | MMNVAVTATPSSLLYSPLLLPSQGPNRRMQWKRNGKRRLGTKVAVSGVITAGFELKPPPYPLDALEPHMSRETLDYHWGKHHKTYVENLNKQILGTDLDALSLEEVVLLSYNKGNMLPAFNNAAQAWNHEFFWESIQPGGGGKPTGELLRLIERDFGSFEEFLERFKSAAASNFGSGWTWLAYKANRLDVANAVNPLPKEEDKKLVIVKTPNAVNPLVWDYSPLLTIDTWEHAYYLDFENRRAEYINTFMEKLVSWETVSTRLESAIARAVQREQEGTETEDEENPDDEVPEVYLDSDIDVSEVD |
| NP_197722.1 | FSD3 Arabidopsis thaliana | MSSCVVTTSCFYTISDSSIRLKSPKLLNLSNQQRRRSLRSRGGLKVEAYYGLKTPPYPLDALEPYMSRRTLEVHWGKHHRGYVDNLNKQLGKDDRLYGYTMEELIKATYNNGNPLPEFNNAAQVYNHDFFWESMQPGGGDTPQKGVLEQIDKDFGSFTNFREKFTNAALTQFGSGWVWLVLKREERRLEVVKTSNAINPLVWDDIPIICVDVWEHSYYLDYKNDRAKYINTFLNHLVSWNAAMSRMARAEAFVNLGEPNIPIA |
| EOY22706.1 | FSD 2 Theobroma cacao | MAAAASMATSLTFPLLPSQGLRGPFTSSLPCTIPQRRFWRKVATNLITAKFELKPPPYTLNALEPHMSRQTLEYHWGKHHRTYVENLNKQIAGTELEGLPLEDIIIVSYNNGDILPAFNNAAQAWNHDFFWESMKPGGGGKPSGDLLDLIERDFGSFEQFIQEFKSAAAAQFGSGWAWLAYKANRLDVENAVNPWPSEKDKKLVVVKSPNAVNPLVWDYFPLLTIDVWEHAYYLDFQNRRPDYISMFMEKLISWEAVSARLEKAKALAAEREMEEERRKKEEEEKQTDGEAVEMYLDSDTDDSEAE |
| XP_010427452.1 | MSD2 Camelina sativa | MTTVVFIIVLVIFAASLYDAGGTTMDPCLESMKTASLPDLPYAYDALEPVISEEIMRLHHQKHHQTYVTQYNKALNSLRSAMADGDHSAVVKLQSLIKFNGGGHVNHAIFWKNLAPVHEGGGKPPHDPLSSAIDAHFGSLEGLIQKMNAEGAAVQGSGWVWLGLDRELKRLVVETTANQDPLVTKGTHLVPLIGIDVWEHAYYPQYKNARAEYLKNIWSVINWKYAADIFEKHSRDLDTN |
| XP_006402995.2 | MSD2 Eutrema salsugineum | MSIVVTIILIFAILAVSLDGKREKTMEPCLESMKTASLPDLPYAYDALEPAISEEIMRLHHQKHHQTYVTQYNKALGSLRSALADGDHSSVVKLQSLIKFNGGGHVNHAIFWKNLAPVHEGGGKPPQDPLCSAIDAHFGSLEGLMQKMNAEGAAVQGSGWVWFGLDKELKRLVVETTANQDPLVTKGPHLVPLIGIDVWEHAYYPQYKNARAEYLKNIWSVINWKYAADIFEKHNRDLNEF |
| XP_013603533.1 | MSD2Brassica oleracea var. oleracea | MSIVVSIILLTVFAMSIDVGTGKSTTMDPCLESMKTASLPDLPYAYDALEPAISEEIMRLHHQKHHQTYVTNYNKALEHLRSALADGDHSSVVKLQSQIKFNGGGHVNHAIFWKNLAPVHEGGGKPPHDPLSSAIDAHFGSLEELMQKMNTEGAAVQGSGWVWFGLDKELKRLVVETTANQDPLVTKGSHLVPLVGIDVWEHAYYPQYKNARAEYLKNIWTVINWKYASDIFVKHNRDLNEF |
| XP_013663290.1 | MSD2 Brassica napus | MIIVVSIILLTVFAMSIDVGTGKSTTMDPCLESMKTASLPDLPYAYDALEPAISEEIMRLHHQKHHQTYVTNYNKALEHLRSALADGDHSSVVKLQSQIKFNGGGHVNHAIFWKNLAPVHEGGGKPPHDPLSSAIDAHFGSLEELMQKMNTEGAAVQGSGWVWFGLDKELKRLVVETTANQDPLVTKGSHLVPLVGIDVWEHAYYPQYKNARAEYLKNIWTVINWKYASDIFVKHNRDLNEF |
| XP_018489893.1 | MSD2 Raphanus sativus | MSIGVYIIFLSVLAMSLDVGRGKTTMDPCLESMKTASLPDLPYPYDALEPAISEEIMRLHHQKHHQTYVTNYNKALDQLRSALANGDHSSVVKLQSQIKFNGGGHVNHAIFWKNLAPVHEGGGKPPHDPLSSAIDAHFGSLEELMQKMNTEGAAVQGSGWVWFGLDKELKRLVVETTANQDPLVTKGSDLVPLIGIDVWEHAYYPQYKNARAEYLKNIWTVINWKYASDIFEKHNRHLNEV |
| XP_009116339.1 | MSD2Brassica rapa | MSIVVSIFLLTVFAMSLDVGTGKTTMEPCLESMKTASLPDLPYAYDALEPAISEEIMRLHHLKHHQTYVTNYNKALDHLRSALSSGDHSSVVKLQSQIKFNGGGHVNHAIFWKNLAPVHEGGGKPPQDPLSSAIDAHFGSLEELMQKMNSEGAAVQGSGWVWFGLDKELKRLVVETTANQDPLVTKGSHLVPLIGIDVWEHAYYPQYKNARAEYLKNIWTVINWKYASDIFEKHNRGLNEF |
| XP_010525403.1 | MSD2 Tarenaya hassleriana | MDPNLQSLKTAVLPELPYPYNALEPAISEEIMRLHHSKHHQTYVNEYNKALEKLRCAMEEGDHSSVVKLHGLIKFNGGGHINHAIFWKNLAPVREGGGQPPTDPLSSSIDTHFGSLEALMQKMNAEGIALQGSGWVWFGLDKELKTLVIETTANQDPLVTKGSNLVPLIGIDVWEHAYYPQYKNARAEYLKNIWSVINWKYAAEVYEREFGL |
| XP_038721861.1 | MSD Tripterygium wilfordii | MALRSLVTRKTLADPRALGLGLAQLRGLKTFSLPDLPYDYGALEPAISGEIMQLHHQKHHQTYITNYNKALEQLQGAMDKGDSPTVVKLQSAIKFNGGGHVNHSIFWKNLTPVHEGGGEPPKGSLGSAIDTHFGSFEALIQKMNAEGAALQGSGWVWLGVDRELKKLLVETTANQDPLVTKGANLVPLLGIDVWEHAYYLQYKNVRPDYLKNIWKVVNWKYASEIFERESQ |
| XP_017650238.1 | MSD Gossypium australe | MALLRSSVTRRTLTLGLNSSRLLPLSRSLQTFSLPDLPYDYGALEPAISGEIMQLHHQKHHQTYITNYNKALEQLHEAIQKGDSSTVVKLQSAIKFNGGGHVNHSIFWKNLAPIREGGGEPPKASLGWAIDSHFGSLESLIQKMNAEGAALQGSGWVWLGVNKELKKLVIETTPNQDPLVTKGPHLVPLLGIDVWEHAYYLQYKNVRPDYLKNIWKVINWKYASEVYEKECA |
| XP_018821863.1 | MSD Juglans regia | MALRSVLTRKTTLGLGLRGIQTFSLPDLPYDYGALEPAISGEIMQLHHHKHHQAYITNYNKALEQLEGALAKGDTTSIVKLQSAIKFNGGGHINHSIFWKNLTPVREGGGEPPHSSLGWAIDTNFGSLEALVQKINSEGAALQGSGWVWFGLDKESKKLVVETTANQDPLVTKGQHLVPLLGIDVWEHAYYLQYKNVRPDYLKNIWKVINWKYASEVYDKESK |
| XP_012456160.1 | MSD Gossypium arboreum | MYKTHISGCTLEKKNLLRLQMALRSLATRKTLTLALNSTRLAQSRALQTFSLPDLPYDYGALEPAISGEIMQLHHQKHHQTYITNYNKALEQLHDAIQKGDSSTVVKLQSAIKFNGGGHINHSIFWKNLAPIREGGGEPPHGSLGWAIDTNFGSLESLIQKMNAEGAALQGSGWVWLGLDKELKKLVVETTANQDPLVTNGPNLVPLLGIDVWEHAYYLQYKNVRPDYLKNVWKVINWKYASEVYEKESA |
| ASL04618.1 | MSD1Actinidia rufa | MALRTLATRRRLGLGFQSIRGFQTFSLPDLPYDYGALEPAISGEIMQLHHQKHHQTYITNYNKALEQLDAAMAKGDAPTVVKLQSAIKFNGGGHVNHSIFWKNLAPVHEGGGEPPKGSLGSTIDTNFGSLEALIQKMNAEGAAVQGSGWVWLGVDKELKKLVIETTANQDPLVTKGSLVPLLGIDVWEHAYYLQYKNVRPDYLKNIWKVMNWKYASEVYEKECP |
| XP_006407489.1 | MSD1 Eutrema salsugineum | MAIRSVATRKTLAGLKETSSRLLGFRGIQTFTLPDLPYDYSALEPAISGEIMQLHHQKHHQTYVTNYNNALEQLDQAVNKGDASTVVKLQSAIKFNGGGHVNHSIFWKNLAPVNQGGGEPPKGALGGAIDTHFGSLEGLVKKMNAEGAALQGSGWVWLGLDKELKKLVVDTTANQDPLVTKGASLVPLVGIDVWEHAYYLQYKNVRPDYLKNVWKVINWKYASEVYEKECK |
| KAA3481249.1 | MSD Gossypium raimondii | MALRSSVTRRTLTLGLNSTRLLPLSRSLQTFSLPDLPYDYGALEPAISGEIMQLHHQKHHQTYITNYNKALEQLHEAIQKGDSSTVVKLQSAIKFNGGGHVNHSIFWKNLAPIREGGGEPPKASLGWAIDTHFGSLESLIQKMNTEGAALQGSGWVWLGVNKELKKLVIETTPNQDPLVTKGPHLVPLLGIDVWEHAYYLQYKNVRPDYLKNIWKVINWKYASEVYEKECA |
| XP_009122529.2 | MSD1Brassica rapa | MAIRSLASRRTLAGLKETSSRLLGLRSIQTFTLPDLPYDYSALEPAISGEIMQIHHQKHHQAYVTNYNNALEQLDQAVNKGDASAVVKLQSAIKFNGGGHVNHSIFWKNLAPVKEGGGEPPKGALGGAIDTHFGSLEGLVKKMSAEGAALQGSGWVWLGLDKELKKLVVDTTANQDPLVTKGGSLVPLVGIDVWEHAYYLQYKNVRPEYLKNVWKVINWKYASEVYEKECK |
| XP_010490355.1 | MSD1 Camelina sativa | MAIRCVASRKTLAGLKETSSRLIGFRGIQTFTLPDLPYDYGALEPAISGEIMQIHHQKHHQTYVTNYNNALEQLDQAVSKGDASAVVKLQSAIKFNGGGHVNHLIFWKNLAPVKEGGGEPPKGSLGSAIDTHFGSLEGLVKKMSAEGAAVQGSGWVWLGLDKELKKLVVDTTANQDPLVTKGGSLVPLVGIDVWEHAYYLQYKNVRPEYLKNVWKVINWKYASEIYEKECN |
| NP_001313781.1 | MSD Gossypium hirsutum | MALRSSITRRTLTLGLNSTMLLPLSRSLQTFSLPDLPYDYGALEPAISGEIMQLHHQKHHQTYITNYNKALEQLHEAIQKGDSSTVVKLQSAIKFNGGGHVNHSIFWKNLAPIREGGGEPPKASLGWAIDTHFGSLESLIQKMNAEGAGLQGSGWVWVGVNKELKKLVIETTPNQDPLVTKGPHLVPLLGIDVWEHAYYLQYKNVRPDYLKNIWKVINWKYASEVYEKECA |
| ABL75952.1 | MSD Eutrema halophilum | MAIRSVATRKTLAGLKETSSRLLGFRGIQTFTLPDLPYDYSALEPAISGEIMQLHHQKHHQTYVTNYNNALEQLDQAVNKGDASTVVKLQSAIKFNGGGHVNHSIFWKNLAPVNQGGGEPPKGALGGAIDTHFGSLEGLVKKMNAEGAALQGSGWVWLGLDKELKKLVVDTTANQDPLVTKGASLVPLVGIDVWEHAYYLQYKNVRPDYLKNVWKVINWKYASEVYEKECN |
| NP_001302584.1 | MSD1 Brassica napus | MAIRSLASRRTLAGLKETSSRLLGLRSIQTFTLPDLPYDYSALEPAISGEIMQIHHQKHHQAYVTNYNNALEQLDQAVNKGDASTVVKLQSAIKFNGGGHVNHSIFWKNLAPVKEGGGEPPKGALGGAIDTHFGSLEGLVKKMSAEGAALQGSGWVWLGLDKELKTLVVDTTANQDPLVTKGGSLVPLVGIDVWEHAYYLQYKNVRPEYLKNVWKVINWKYASEVYEKECK |
| ASL04618.1 | MSD Actinidia deliciosa | MALRTLATRRRLGLGFQSIRGLQTFSLPDLPYDYGALEPAISGEIMQLHHQKHHQTYITNYNKALEQLDAAMAKGDAPTVVKLQSAIKFNGGGHVNHSIFWKNLAPVREGGGEPPKGSLGSTIDTNFGSLEALIQKMNAEGAAVQGSGWVWLGVDKELKKLVIETTANQDPLVTKGSLVPLLGIDVWEHAYYLQYKNVRPDYLKNIWKVMNWKYASQVYEKECP |
| XP_018433955.1 | MSD1Raphanus sativus | MAIRSVASRRTLAGLKETSSRLLGSRSIQTFTLPDLPYDYSALEPAISGEIMQIHHQKHHQAYVTNYNNALEQLDQAVNKGDASAVVKLQSAIKFNGGGHVNHSIFWKNLAPVKEGGGEPPKGSLGGAIDTHFGSLEGLVKKMSAEGAAVQGSGWVWLGLDKELKKLVVDTTANQDPLVTKGGSLVPLVGIDVWEHAYYLQYKNVRPDYLKNVWKVINWKYASEVYEKECK |
| NP_187703.1 | MSD1Arabidopsis thaliana | MAIRCVASRKTLAGLKETSSRLLRIRGIQTFTLPDLPYDYGALEPAISGEIMQIHHQKHHQAYVTNYNNALEQLDQAVNKGDASTVVKLQSAIKFNGGGHVNHSIFWKNLAPSSEGGGEPPKGSLGSAIDAHFGSLEGLVKKMSAEGAAVQGSGWVWLGLDKELKKLVVDTTANQDPLVTKGGSLVPLVGIDVWEHAYYLQYKNVRPEYLKNVWKVINWKYASEVYEKENN |
| NP_191194.1 | MSD2 Arabidopsis thaliana | MTTTVIIIIFVAIFATTLHDARGATMEPCLESMKTASLPDLPYAYDALEPAISEEIMRLHHQKHHQTYVTQYNKALNSLRSAMADGDHSSVVKLQSLIKFNGGGHVNHAIFWKNLAPVHEGGGKPPHDPLASAIDAHFGSLEGLIQKMNAEGAAVQGSGWVWFGLDRELKRLVVETTANQDPLVTKGSHLVPLIGIDVWEHAYYPQYKNARAEYLKNIWTVINWKYAADVFEKHTRDLDIN |
| RWR80468.1 | MSD Cinnamomum micranthum f. kanehirae | MALRSLLSKRTLGLGFGQVRGLVTFSLPDLPYDYSALEPAISGEIMQLHHQKHHQTYVTNYNKALEQLDDAMAKGDAPTVVKLQSAIKFNGGGHINHSIFWKNLIPIHEGGGEPPKSSLGWAIDMHFGSFEALLQKMNAEGAALQGSGWVWLGLDKELKRLVVETTANQDPLVTKGASLVPLLGIDVWEHAYYLQYKNVRPDYLNNIWKVINWKYASELYEKESP |
| XP_021618226.1 | MSD Manihot esculenta | MALRSLVTRKNLSSAFKAATGLGQLRGLQTFSLPDLPYDYGALEPAISGEIMQLHHQKHHQTYITNYNKALEQLNDAMEKGDSATVVKLQSAIKFNGGGHVNHSIFWKNLSPVREGGGEPPHGSLGWAIDADFGSLEKLIQKMNAEGAAVQGSGWVWLAVDKELKKLVVETTANQDPLVTKGPTLVPLLGIDVWEHAYYLQYKNVRPDYLKNIWKVMNWKYASEVYAKECPSS |
| OVA03276.1 | MSD Macleaya cordata | MALRTVLSRKTLGLGLGFQNARLGFSNVRELQTFTLPDLPYDYGALEPAISGEIMQLHHQKHHQTYITNYNKALEQLHEAMVKGDASTVVKLQGAIKFNGGGHVNHSIFWKNLTPVREGGGEPPKGSLASAIDTHFGSFESLVQKINGEGAALQGSGWVWLGLDKELKKLVVETTSNQDPLVTKGPNLVPLLGIDVWEHAYYLQYKNVRPDYLKNIWKVINWKYAGEVYEKEFP |
| QCD25905.1 | MSD1 Helianthus tuberosus | MALRALTNVKTLGRLRHQQIRGLQTFTLPDLSYDYGALEPAISGEIMQLHHQKHHQTYITNYNKALEQLDDAIAKGDASTAVKLQSAIKFNGGGHVNHSIFWKNLAPTHEGGGEPPHGSLGWAIDQHFGSMEKLVAKMNAEGAAVQGSGWVWLAVDKELKRLVVETTANQDPLVTKGASLVPLVGIDVWEHAYYLQYKNVRPDYLKNIWKVINWKYASEVYEKECP |
| XP_007011340.1 | MSD Theobroma cacao | MALRSLATRKTLTLGLNSTRLTQSRALQTFSLPDLPYDYGALEPAISGEIMQLHHQKHHQTYITNYNKALEQLHEAIQKGDSSTVVKLQSAIKFNGGGHINHSIFWKNLAPIHEGGGEPPKGSLGWAIDTSFGSLESLIQKMNAEGAALQGSGWVWLGVDKELKKLVIETTANQDPLVTKGPALVPLLGIDVWEHAYYLQYKNVRPDYLKNIWKVIDWKYASEVYEKECP |
| XP_013617540.1 | MSD1Brassica oleracea var. oleracea | MAIRSLAGRRTLTGLKETSSRLLGFRSIQTFTLPDLPYDYSALEPAISGEIMQIHHQKHHQAYVTNYNNALEQLDQAVNKGDASGVVKLQSAIKFNGGGHVNHSIFWKNLAPVKEGGGEPPKGALGYAIDTHFGSLEGLVKKMSAEGAALQGSGWVWLGLDKELKKLVVDTTANQDPLVTKGGSLVPLVGIDVWEHAYYLQYKNVRPEYLKNVWKVINWKYASEVYEKECK |
| XP_034705342.1 | MSD Vitis riparia | MALRTLITRRSLGLGLGVSQSVRGLQTVSLPDLPYDYGALEPAISGEIMKLHHQKHHQTYITNYNKALEQLHEAMEKGDSSTVVKLQGAIKFNGGGHVNHSIFWKNLTPVHEGGGEPPKGSLGWAIDTHFGSMEALVAKINSEGAAVQGSGWVWLGLDKQLKKLVVETTANQDPLVTKGPNLVPLLGIDVWEHAYYLQYKNVRPDYLKNIWKVIDWKYASEVYEKECP |
| OMO63671.1 | MSD Corchorus capsularis | MALRCLATRKTLAQAALNSTRLTQSRALQTFSLPDLPYDYGALEPAISGEIMQLHHQKHHQTYITNYNKALEQLHEAIEKGDASTVVKLQSAIKFNGGGHVNHSIFWKNLAPVREGGGEPPHGSLGWAIDTNFGSLESLIQKMNAEGAALQGSGWVWLAVDKELKKLVIETTANQDPLVTKGASLVPLLGIDVWEHAYYLQYKNVRPDYLKNIWKVINWKYASSVYENEFPSARSP |
| OMO89989.1 | MSD Corchorus olitorius | MALRCLATRKTLAQAAFNSTRLTQSRALQTFSLPDLPYDYGALEPAISGEIMQLHHQKHHQTYITNYNKALEQLHEAIEKGDASTVVKLQSAIKFNGGGHVNHSIFWKNLAPVREGGGEPPHGSLGWAIDTNFGSLESLIQKMNAEGAALQGSGWVWLAVDKELKKLVIETTANQDPLVTKGASLVPLLGIDVWEHAYYLQYKNVRPDYLKNIWEVINWKYASSVYENEFPSARSP |
| XP_017422543.1 | MSD Vigna angularis | MAARALLTRKTLASVLRNDARPLGVGAAVATHSRGLHVYTLPDLDYDYGALEPAISGEIMQLHHQKHHQTYITNYNKALEQLQDAVAKADSSAVVKLQAAIKFNGGGHINHSIFWKNLAPVREGGGEPPKGPLGWAIDTHFGSFEALIQKVNAEGAALQGSGWVWLGLDKELKRLVVETTANQDPLVTKGPNLVPLLGIDVWEHAYYLQYKNVRPDYLKNIWKVINWKYANDVYEKESS |
| XP_010670629.1 | MSD Beta vulgaris subsp. vulgaris | MALRVLMTRKTLASAKQGLLQCRSLQTFSLPDLPYDYGALQPAISGEIMQIHHQKHHQTYITNYNKALEQLDDAIAKGDASSVVKLQSAIKFNGGGHVNHSIFWKNLAPINEGGGEPPKSSLGWAIDSNFGSLEALIQKMNAEGAAVQGSGWVWLGLDTQSKKLVVETTPNQDPLVTKGPSLVPILGIDVWEHAYYLQYKNVRPDYLKNIWKVINWKYASEVYEKEVPSH |
| XP_027110626.1 | MSD Coffea arabica | MALRTLVTRKALGNSVAFRQQLRGLQTYTLPDLPYDYGAIEPAISGEIMQLHHQKHHQTYVTNFNKALEQLDDAINKGDAPTVVKLQSAIKFNGGGHINHSIFWKNLAPIREGGGEPPKGSLGWAIDNHFGSLEALVQKMNADGAGLQGSGWVWLGLDKELKRLVVETTANQDPLVTKGSSLVPLLGIDVWEHAYYLQYKNVRPDYLKNIWKVINWKYASDIYEKECP |
| KAB1217601.1 | MSD Morella rubra | MALRPLLTRKTLGLRLPQLRGLQTFSLPDLPYDYGALEPVISGEIMQLHHQKHHQAYITNFNKALEQLDEAMAKGDSTAVVKLQSAIKFNGGGHINHSIFWKNLAPIREGGGEPPKGSLGWAIDTHFGSLETLVQKMNLEGAALQGSGWVWLGLDKELKKLVVETTANQDPLVTKGPHLVPLLGIDVWEHAYYLQYKNVRPDYLKNVWKVIDWHYSSEVYEKESK |
| XP_031372915.1 | MSD Punica granatum | MALRTLLSCRTLAAAVKPELGFSVGLRSLVTFSLPDLPYDYGALEPAISGEIMQLHHQKHHQAYITNYNKALEQLELAMSKGDAPSVVKLQSAIKFNGGGHVNHSIFWKNLAPVREGGGEPPKGSLGWAIDTHFGSLEALAQKMSTEGAALQGSGWVWLGVDKELKKLVVETTVNQDPLVTKGPSLVPLVGIDVWEHAYYLQYKNVRPDYLKNIWKVINWKYASEVFEKECP |
| XP_020887352.1 | MSD1 Arabidopsis lyrata subsp. lyrata | MAIRCVASRKTLAGLKETSSRLLGFRGVQTFTLPDLPYDYGALEPAISGEIMQIHHQKHHQAYVTNYNNALEQLDQAVNKGDASTVVKLQSAIKFNGGGHVNHSIFWKNLAPVNEGGGEPPKGSLGSAIDTHFGSPEGLVKKMSAEGAAVQGSGWVWLGLDKELKKLVVDTTANQDPLVTKGGSLVPLVGIDVWEHAYYLQYKNVRPEYLKNVWKVINWKYASEVYEKECN |

**Table S2**. Predicted PTM sites of MSD2.

| **ID** | **Position** | **Residue** | **PTM scores** | **Cutoff=0.5** |
| --- | --- | --- | --- | --- |
| sp\|Q9LYK8\|SODM2_ARATH | 2 | T | Phosphothreonine:0.068;O-linked_glycosylation:0.107 | None |
| sp\|Q9LYK8\|SODM2_ARATH | 3 | T | Phosphothreonine:0.066;O-linked_glycosylation:0.165 | None |
| sp\|Q9LYK8\|SODM2_ARATH | 4 | T | Phosphothreonine:0.081;O-linked_glycosylation:0.096 | None |
| sp\|Q9LYK8\|SODM2_ARATH | 16 | T | Phosphothreonine:0.045;O-linked_glycosylation:0.062 | None |
| sp\|Q9LYK8\|SODM2_ARATH | 17 | T | Phosphothreonine:0.048;O-linked_glycosylation:0.066 | None |
| sp\|Q9LYK8\|SODM2_ARATH | 22 | R | Methylarginine:0.069 | None |
| sp\|Q9LYK8\|SODM2_ARATH | 25 | T | Phosphothreonine:0.101;O-linked_glycosylation:0.403 | None |
| sp\|Q9LYK8\|SODM2_ARATH | 28 | P | Hydroxyproline:0.086 | None |
| sp\|Q9LYK8\|SODM2_ARATH | 29 | C | S-palmitoyl_cysteine:0.034 | None |
| sp\|Q9LYK8\|SODM2_ARATH | 32 | S | Phosphoserine:0.184;O-linked_glycosylation:0.224 | None |
| sp\|Q9LYK8\|SODM2_ARATH | 34 | K | Ubiquitination:0.381;SUMOylation:0.05;N6-acetyllysine:0.137;Methyllysine:0.172;Hydroxylysine:0.035 | None |
| sp\|Q9LYK8\|SODM2_ARATH | 35 | T | Phosphothreonine:0.155;O-linked_glycosylation:0.498 | None |
| sp\|Q9LYK8\|SODM2_ARATH | 37 | S | Phosphoserine:0.682;O-linked_glycosylation:0.269 | Phosphoserine:0.682 |
| sp\|Q9LYK8\|SODM2_ARATH | 39 | P | Hydroxyproline:0.037 | None |
| sp\|Q9LYK8\|SODM2_ARATH | 42 | P | Hydroxyproline:0.047 | None |
| sp\|Q9LYK8\|SODM2_ARATH | 43 | Y | Phosphotyrosine:0.107 | None |
| sp\|Q9LYK8\|SODM2_ARATH | 45 | Y | Phosphotyrosine:0.068 | None |
| sp\|Q9LYK8\|SODM2_ARATH | 50 | P | Hydroxyproline:0.135 | None |
| sp\|Q9LYK8\|SODM2_ARATH | 53 | S | Phosphoserine:0.129;O-linked_glycosylation:0.082 | None |
| sp\|Q9LYK8\|SODM2_ARATH | 58 | R | Methylarginine:0.097 | None |
| sp\|Q9LYK8\|SODM2_ARATH | 62 | Q | Pyrrolidone_carboxylic_acid:0.065 | None |
| sp\|Q9LYK8\|SODM2_ARATH | 63 | K | Ubiquitination:0.199;SUMOylation:0.04;N6-acetyllysine:0.315;Methyllysine:0.08;Hydroxylysine:0.031 | None |
| sp\|Q9LYK8\|SODM2_ARATH | 66 | Q | Pyrrolidone_carboxylic_acid:0.042 | None |
| sp\|Q9LYK8\|SODM2_ARATH | 67 | T | Phosphothreonine:0.126;O-linked_glycosylation:0.056 | None |
| sp\|Q9LYK8\|SODM2_ARATH | 68 | Y | Phosphotyrosine:0.13 | None |
| sp\|Q9LYK8\|SODM2_ARATH | 70 | T | Phosphothreonine:0.106;O-linked_glycosylation:0.047 | None |
| sp\|Q9LYK8\|SODM2_ARATH | 71 | Q | Pyrrolidone_carboxylic_acid:0.035 | None |
| sp\|Q9LYK8\|SODM2_ARATH | 72 | Y | Phosphotyrosine:0.061 | None |
| sp\|Q9LYK8\|SODM2_ARATH | 73 | N | N-linked_glycosylation:0.036 | None |
| sp\|Q9LYK8\|SODM2_ARATH | 74 | K | Ubiquitination:0.347;SUMOylation:0.046;N6-acetyllysine:0.401;Methyllysine:0.17;Hydroxylysine:0.093 | None |
| sp\|Q9LYK8\|SODM2_ARATH | 77 | N | N-linked_glycosylation:0.035 | None |
| sp\|Q9LYK8\|SODM2_ARATH | 78 | S | Phosphoserine:0.247;O-linked_glycosylation:0.055 | None |
| sp\|Q9LYK8\|SODM2_ARATH | 80 | R | Methylarginine:0.072 | None |
| sp\|Q9LYK8\|SODM2_ARATH | 81 | S | Phosphoserine:0.208;O-linked_glycosylation:0.052 | None |
| sp\|Q9LYK8\|SODM2_ARATH | 89 | S | Phosphoserine:0.229;O-linked_glycosylation:0.077 | None |
| sp\|Q9LYK8\|SODM2_ARATH | 90 | S | Phosphoserine:0.539;O-linked_glycosylation:0.051 | Phosphoserine:0.539 |
| sp\|Q9LYK8\|SODM2_ARATH | 93 | K | Ubiquitination:0.3;SUMOylation:0.042;N6-acetyllysine:0.22;Methyllysine:0.1;Hydroxylysine:0.028 | None |
| sp\|Q9LYK8\|SODM2_ARATH | 95 | Q | Pyrrolidone_carboxylic_acid:0.128 | None |
| sp\|Q9LYK8\|SODM2_ARATH | 96 | S | Phosphoserine:0.229;O-linked_glycosylation:0.059 | None |
| sp\|Q9LYK8\|SODM2_ARATH | 99 | K | Ubiquitination:0.62;SUMOylation:0.043;N6-acetyllysine:0.279;Methyllysine:0.137;Hydroxylysine:0.063 | Ubiquitination:0.62 |
| sp\|Q9LYK8\|SODM2_ARATH | 101 | N | N-linked_glycosylation:0.032 | None |
| sp\|Q9LYK8\|SODM2_ARATH | 107 | N | N-linked_glycosylation:0.033 | None |
| sp\|Q9LYK8\|SODM2_ARATH | 113 | K | Ubiquitination:0.37;SUMOylation:0.045;N6-acetyllysine:0.188;Methyllysine:0.229;Hydroxylysine:0.101 | None |
| sp\|Q9LYK8\|SODM2_ARATH | 114 | N | N-linked_glycosylation:0.034 | None |
| sp\|Q9LYK8\|SODM2_ARATH | 117 | P | Hydroxyproline:0.116 | None |
| sp\|Q9LYK8\|SODM2_ARATH | 124 | K | Ubiquitination:0.507;SUMOylation:0.138;N6-acetyllysine:0.146;Methyllysine:0.119;Hydroxylysine:0.074 | Ubiquitination:0.507 |
| sp\|Q9LYK8\|SODM2_ARATH | 125 | P | Hydroxyproline:0.101 | None |
| sp\|Q9LYK8\|SODM2_ARATH | 126 | P | Hydroxyproline:0.112 | None |
| sp\|Q9LYK8\|SODM2_ARATH | 129 | P | Hydroxyproline:0.108 | None |
| sp\|Q9LYK8\|SODM2_ARATH | 132 | S | Phosphoserine:0.177;O-linked_glycosylation:0.078 | None |
| sp\|Q9LYK8\|SODM2_ARATH | 140 | S | Phosphoserine:0.675;O-linked_glycosylation:0.083 | Phosphoserine:0.675 |
| sp\|Q9LYK8\|SODM2_ARATH | 146 | Q | Pyrrolidone_carboxylic_acid:0.058 | None |
| sp\|Q9LYK8\|SODM2_ARATH | 147 | K | Ubiquitination:0.289;SUMOylation:0.046;N6-acetyllysine:0.179;Methyllysine:0.125;Hydroxylysine:0.043 | None |
| sp\|Q9LYK8\|SODM2_ARATH | 149 | N | N-linked_glycosylation:0.033 | None |
| sp\|Q9LYK8\|SODM2_ARATH | 156 | Q | Pyrrolidone_carboxylic_acid:0.115 | None |
| sp\|Q9LYK8\|SODM2_ARATH | 158 | S | Phosphoserine:0.063;O-linked_glycosylation:0.244 | None |
| sp\|Q9LYK8\|SODM2_ARATH | 167 | R | Methylarginine:0.017 | None |
| sp\|Q9LYK8\|SODM2_ARATH | 170 | K | Ubiquitination:0.273;SUMOylation:0.085;N6-acetyllysine:0.207;Methyllysine:0.165;Hydroxylysine:0.06 | None |
| sp\|Q9LYK8\|SODM2_ARATH | 171 | R | Methylarginine:0.015 | None |
| sp\|Q9LYK8\|SODM2_ARATH | 176 | T | Phosphothreonine:0.113;O-linked_glycosylation:0.316 | None |
| sp\|Q9LYK8\|SODM2_ARATH | 177 | T | Phosphothreonine:0.116;O-linked_glycosylation:0.246 | None |
| sp\|Q9LYK8\|SODM2_ARATH | 179 | N | N-linked_glycosylation:0.034 | None |
| sp\|Q9LYK8\|SODM2_ARATH | 180 | Q | Pyrrolidone_carboxylic_acid:0.127 | None |
| sp\|Q9LYK8\|SODM2_ARATH | 182 | P | Hydroxyproline:0.05 | None |
| sp\|Q9LYK8\|SODM2_ARATH | 185 | T | Phosphothreonine:0.092;O-linked_glycosylation:0.27 | None |
| sp\|Q9LYK8\|SODM2_ARATH | 186 | K | Ubiquitination:0.33;SUMOylation:0.116;N6-acetyllysine:0.132;Methyllysine:0.16;Hydroxylysine:0.023 | None |
| sp\|Q9LYK8\|SODM2_ARATH | 188 | S | Phosphoserine:0.072;O-linked_glycosylation:0.237 | None |
| sp\|Q9LYK8\|SODM2_ARATH | 192 | P | Hydroxyproline:0.099 | None |
| sp\|Q9LYK8\|SODM2_ARATH | 203 | Y | Phosphotyrosine:0.054 | None |
| sp\|Q9LYK8\|SODM2_ARATH | 204 | Y | Phosphotyrosine:0.051 | None |
| sp\|Q9LYK8\|SODM2_ARATH | 205 | P | Hydroxyproline:0.068 | None |
| sp\|Q9LYK8\|SODM2_ARATH | 206 | Q | Pyrrolidone_carboxylic_acid:0.25 | None |
| sp\|Q9LYK8\|SODM2_ARATH | 207 | Y | Phosphotyrosine:0.055 | None |
| sp\|Q9LYK8\|SODM2_ARATH | 208 | K | Ubiquitination:0.427;SUMOylation:0.033;N6-acetyllysine:0.171;Methyllysine:0.545;Hydroxylysine:0.089 | Methyllysine:0.545 |
| sp\|Q9LYK8\|SODM2_ARATH | 209 | N | N-linked_glycosylation:0.033 | None |
| sp\|Q9LYK8\|SODM2_ARATH | 211 | R | Methylarginine:0.049 | None |
| sp\|Q9LYK8\|SODM2_ARATH | 214 | Y | Phosphotyrosine:0.091 | None |
| sp\|Q9LYK8\|SODM2_ARATH | 216 | K | Ubiquitination:0.654;SUMOylation:0.031;N6-acetyllysine:0.775;Methyllysine:0.293;Hydroxylysine:0.021 | Ubiquitination:0.654\|N6-acetyllysine:0.775 |
| sp\|Q9LYK8\|SODM2_ARATH | 217 | N | N-linked_glycosylation:0.036 | None |
| sp\|Q9LYK8\|SODM2_ARATH | 220 | T | Phosphothreonine:0.08;O-linked_glycosylation:0.049 | None |
| sp\|Q9LYK8\|SODM2_ARATH | 223 | N | N-linked_glycosylation:0.034 | None |
| sp\|Q9LYK8\|SODM2_ARATH | 225 | K | Ubiquitination:0.349;SUMOylation:0.044;N6-acetyllysine:0.257;Methyllysine:0.082;Hydroxylysine:0.024 | None |
| sp\|Q9LYK8\|SODM2_ARATH | 226 | Y | Phosphotyrosine:0.057 | None |
| sp\|Q9LYK8\|SODM2_ARATH | 233 | K | Ubiquitination:0.611;SUMOylation:0.104;N6-acetyllysine:0.727;Methyllysine:0.131;Hydroxylysine:0.048 | Ubiquitination:0.611\|N6-acetyllysine:0.727 |
| sp\|Q9LYK8\|SODM2_ARATH | 235 | T | Phosphothreonine:0.131;O-linked_glycosylation:0.111 | None |
| sp\|Q9LYK8\|SODM2_ARATH | 236 | R | Methylarginine:0.016 | None |
| sp\|Q9LYK8\|SODM2_ARATH | 241 | N | N-linked_glycosylation:0.048 | None |

**Table S3.** Tyr-nitration prediction by GPS-YNO2.

| Position | Peptide | Score | Cutoff |
| --- | --- | --- | --- |
| 43 | KTASLPDLPYAYDA | 0.532 | 0 |
| 45 | ASLPDLPYAYDALE | 0.375 | 0 |
| 68 | QKHHQTYVTQYNK | 2.465 | 0 |
| 72 | QTYVTQYNKALNS | 1.066 | 0 |
| 203 | LIGIDVWEHAYYP | 0.212 | 0 |
| 204 | GIDVWEHAYYPQ | 0.171 | 0 |
| 207 | HAYYPQYKNARA | 0.217 | 0 |
| 214 | NARAEYLKNIWT | 0.581 | 0 |
| 226 | INWKYAADVFE | 0.475 | 0 |

**Table S4.** List of primers used in this study.

| Name | Sequences |
| --- | --- |
| MSD2 CDS_F | ATGACGACCACCGTTATTATC |
| MSD2 CDS_R | TCAGTTAATATCAAGATCACGA |
| MSD2 promoter_F | CACACAATCCAATGGCAATGATG |
| MSD2 promoter_R | ATCTTTTTATATATGTAGACA |
| MSD2/Y68F_F | ATCAGAAACACCACCAGACTTTTGTCACTCAGTAC |
| MSD2/Y68F_R | AAAGTCTGGTGGTGTTTCTGATGGTGTAGCC |
| qMSD2_F | TGCCATCTTCGCTACGACTC |
| qMSD2_R | AGAACGAAGGCTGTTGAGGG |
| qUBQ5_F | ACCCCTTGAGGTTGAATCATC |
| q UBQ5_R | GTCCTTCTTTCTGGTAAACGT |
